# SINEUP long non-coding RNA acts via PTBP1 and HNRNPK to promote translational initiation assemblies

**DOI:** 10.1101/664029

**Authors:** Naoko Toki, Hazuki Takahashi, Silvia Zucchelli, Stefano Gustincich, Piero Carninci

## Abstract

SINEUPs are long non-coding RNAs (lncRNAs) that contain a SINE element, which up-regulate the translation of target mRNA and have been studied in a wide range of applications for biological and therapeutic tools, although the molecular mechanism is unclear. Here, we focused on the kinetic distribution of target mRNAs and *SINEUP* RNAs by performing co-transfection of expression vectors for these transcripts into human embryonic normal kidney cells (HEK293T/17) to investigate the network of translational regulation. The results showed that co-localization of target mRNAs and *SINEUP* RNAs in the cytoplasm was one of the key phenomena. We identified PTBP1 and HNRNPK as essential RNA binding proteins. These proteins contributed to *SINEUP* RNA sub-cellular distribution and to assembly of translational initiation complexes, leading to enhanced target mRNA translation. These findings will promote a better understanding of the mechanisms on the fate of regulatory RNAs implicated in efficient protein translation.

## Introduction

More than 70% of the mammalian genome is transcribed into RNAs (Carninci *et al*, 2005; The ENCODE Project Consortium, 2012). The majority of these RNAs do not encode proteins and are called non-coding RNAs. A large fraction of non-coding RNAs are found as sense-antisense pairwise transcripts that are co-regulated at the transcription level (Katayama *et al*, 2005). Although some of these non-coding RNAs, such as transfer RNA (tRNA) and ribosomal RNA (rRNA), have been intensely studied, the biological function of 98.5% of the long non-coding RNAs (lncRNAs), which are defined as non-coding RNAs longer than 200 nucleotides (Kapranov *et al*, 2007), are still unknown (de Hoon *et al*, 2015). Importantly, some lncRNAs have regulatory functions at the transcriptional or translational levels (Hu *et al*, 2012; Rinn & Chang, 2012; Quinn & Chang, 2016).

We previously found AS-*Uchl1*, a natural antisense (AS) lncRNA, was able to enhance the translation of its sense *Uchl1* (ubiquitin carboxyterminal hydrolase L1) mRNA in mouse dopaminergic cells (Carrieri *et al* 2012). The functional element embedded in AS-*Uchl1* is an inverted SINEB2 (Short Interspersed Nuclear Element B2) that acts as an effector domain (ED) and is essential for up-regulating protein synthesis (Zucchelli *et al*, 2015). This element enhanced the UCHL1 protein level without changing the *Uchl1* mRNA level. In mouse dopaminergic neurons, mature *Uchl1* mRNA is predominantly localized in the cytoplasm, whereas the natural AS-*Uchl1* RNA is localized in the nucleus at steady state levels. Interestingly, in rapamycin-induced stress conditions, endogenous AS-*Uchl1* RNA is transported to the cytoplasm to increase *Uchl1* mRNA association to heavy polysomes and enhances its translation (Carrieri *et al*, 2012).

We also found that other natural antisense lncRNAs containing SINEs specifically up-regulate translation of the overlapping sense mRNAs, and termed such functional class of lncRNAs “SINEUPs” (Zucchelli *et al*, 2015). In addition to the inverted SINEB2, SINEUPs also require an overlapping region known as the target binding domain (BD) of the sense mRNA (Zucchelli *et al*, 2015). We used the configuration of the original natural SINEUPs to produce a synthetic SINEUP targeting green fluorescent protein (*SINEUP-GFP*) by replacing the BD of AS-*Uchl1* with that of the target enhanced green fluorescent protein (EGFP) mRNA. *SINEUP-GFP* successfully up-regulated the translation of *EGFP* mRNA in an ED-dependent fashion maintaining target mRNA levels unchanged. Despite the functional evidence, the mechanism by which natural and synthetic SINEUPs enhance the translation of a target mRNA is still unknown; in particular, the fate of *SINEUP* RNAs after transcription, such as their subcellular distribution and contribution to protein translation, is unclear.

Here we show that *SINEUP* RNA interacts with nucleocytoplasmic proteins such as PTBP1 (polypyrimidine tract binding protein-1) and HNRNPK (heterogeneous nuclear ribonucleoprotein K) that are essential for RNA localization and translational initiation assembly.

## Results

### SINEUP-GFP enhances *EGFP* mRNA translation

To confirm translational up-regulation activity by SINEUP constructs, we produced synthetic SINEUPs targeting *EGFP* mRNA (Fig 1A). We previously reported that *SINEUP-GFP* enhances EGFP levels (Indrieri *et al*, 2016) more efficiently when cloned into the pCS2+ plasmid than when cloned into pcDNA3.1 plasmid as shown in our previous studies (Carrieri *et al*, 2012, Zucchelli *et al*, 2015). The expression levels of RNAs produced by different plasmids often differ due to different promoters, stability, and polyadenylation status, and in the current study we found that the level of *SINEUP* RNA transcripts was higher for pCS2+ than for pcDNA3.1 (Fig EV1). Therefore, pCS2+ was used for all subsequent experiments. We then examined the EGFP up-regulation activities of SINEUP-GFP and of the deletion mutants constructed in this study; the BD mutant (SINEUP-SCR), the ED deletion mutant (SINEUP-ΔSB2), and the Alu element deletion mutant (SINEUP-ΔAlu) in pCS2+ (Fig 1A). Consistent with previous studies, SINEUP-GFP in pCS2+ showed approximately 2-fold induction of EGFP level compared to the No-insert control (vector containing no SINEUP construct) (Fig 1B, C). SINEUP-SCR and SINEUP-ΔSB2 did not significantly elevate the EGFP level, but the Alu element deletion mutant (SINEUP-ΔAlu) and SINEUP-GFP did significantly elevate the EGFP level. Because none of the constructs significantly affected the *EGFP* mRNA level (Fig 1D), the results indicate that translation of *EGFP* mRNA was induced by SINEUP-GFP and SINEUP-ΔAlu, but not by SINEUP-SCR or SINEUP-ΔSB2.

**Figure 1.**
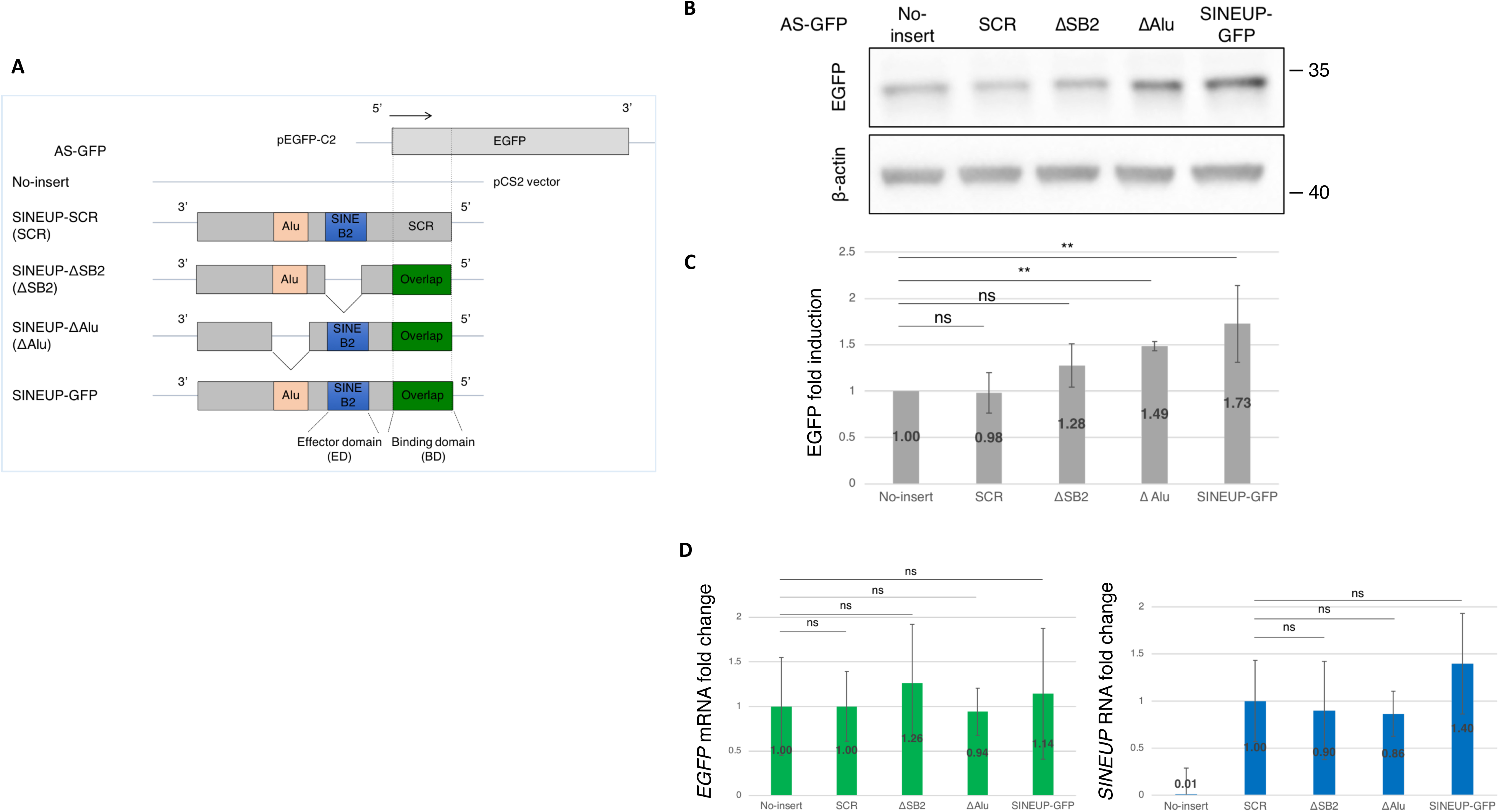
Enhancement of EGFP level by synthetic SINEUP-GFP. A Schematic representation of the SINEUP constructs used in this study. SINEUP-GFP contains the overlap region with *EGFP* (binding domain, BD) and SINEB2 element (effector domain, ED). Domain deletion mutants constructed from SINEUP-GFP are shown: SINEUP-SCR (SCR) contains scrambled sequence instead of *EGFP* BD; SINEUP-delta SB2 (ΔSB2) has a deleted SINEB2 element; and SINEUP-delta Alu (ΔAlu) has a deleted Alu repeat element. B Translational up-regulation of EGFP by co-transfection of EGFP and SINEUP expression vectors. Western blot image showing that the effect of SINEUPs on the EGFP level; the result shown is representative of at least three independent experiments. C Quantification of the up-regulation of EGFP by co-transfection with EGFP and SINEUP vectors. **p < 0.01, ns: not significant, by Student’s *t*-test. Data are means ± SD from at least 3 independent experiments. D Quantification of the *EGFP* mRNA levels following co-transfection with EGFP and SINEUP expression vectors. Data are means ± SD from at least 3 independent experiments.

### *SINEUP* RNAs co-localized with *EGFP* mRNAs in the cytoplasm

We previously showed that, in mouse dopaminergic neurons, mature *Uchl1* mRNA is predominantly localized in the cytoplasm, whereas the natural *SINEUP* RNA AS-*Uchl1* is localized in the nucleus at steady state levels but upon rapamycin treatment it is transported to the cytoplasm to enhance *Uchl1* mRNA translation (Carrieri *et al*, 2012). We hypothesize that the subcellular distribution of *SINEUP* RNAs has a key role in regulating target mRNA translation. To elucidate the kinetic distribution of *EGFP* mRNA and *SINEUP* RNA, we performed RNA FISH (fluorescence *in situ* hybridization) following co-transfection of EGFP and SINEUP expression vectors (SINEUP-GFP or the deletion mutants) into HEK293T/17 cells. We observed that *EGFP* mRNAs were predominantly localized in the cytoplasm (Fig 2A, d, i, n, s), whereas the *SINEUP* RNAs were distributed both in the nucleus and the cytoplasm (Fig 2A, c, h, m, r). In the nucleus, *SINEUP* RNAs were located throughout the nucleoplasm, but not in the nucleolus. For an unknown reason, *SINEUP* RNAs formed intensively clustered spots. In the cytoplasm, *SINEUP* RNAs co-localized with *EGFP* mRNAs, appearing as numerous small dots distributed throughout the cytoplasm (Fig 2A, u, arrows). Co-localization of *EGFP* mRNA and *SINEUP* RNA in the cytoplasm was observed more frequently for SINEUP-GFP (37.75%) and SINEUP-ΔAlu RNAs (30.73%) than for *SINEUP* RNAs with impaired translational up-regulation activity (i.e., SINEUP-SCR and SINEUP-ΔSB2) (Fig 2B); SINEUP-ΔAlu and SINEUP-GFP showed more frequent overlap peaks (arrows) compared with SINEUP-SCR and SINEUP-ΔSB2 (Fig EV2A). This suggests that BD and ED domains contribute both for the translational up-regulation activity and for the co-localization of *EGFP* mRNAs and *SINEUP* RNAs. When the EGFP expression vector was transfected alone, most *EGFP* mRNA was distributed in the cytoplasm (Fig EV2B), as was observed for co-transfection. In contrast, when the expression vectors for *SINEUP* RNAs were transfected alone, the *SINEUP* RNAs were preferably distributed in the nucleus (Fig 2C, e, h, k). To compare the subcellular distribution of *SINEUP* RNAs between cells co-transfected with EGFP and SINEUP vectors to those transfected with SINEUP alone, the signals were detected from whole cell region and the nucleus region by using icy Spot Detector (Fig EV2C). The percentage of *SINEUP-GFP* RNA localized in the nucleus was 60.6% when the SINEUP-GFP vector was transfected alone (Fig 2D); this was significantly reduced to 49.3% and shifted to the cytoplasm when the SINEUP vector was co-transfected with the EGFP vector. A similar finding was observed for SINEUP-ΔAlu, but no significant differences were observed for SINEUP-SCR or SINEUP-ΔSB2 (Fig EV3 A-C). These observations suggest that the *SINEUP* RNAs with translational up-regulation activity were exported to the cytoplasm in the presence of *EGFP* mRNA.

**Figure 2.**
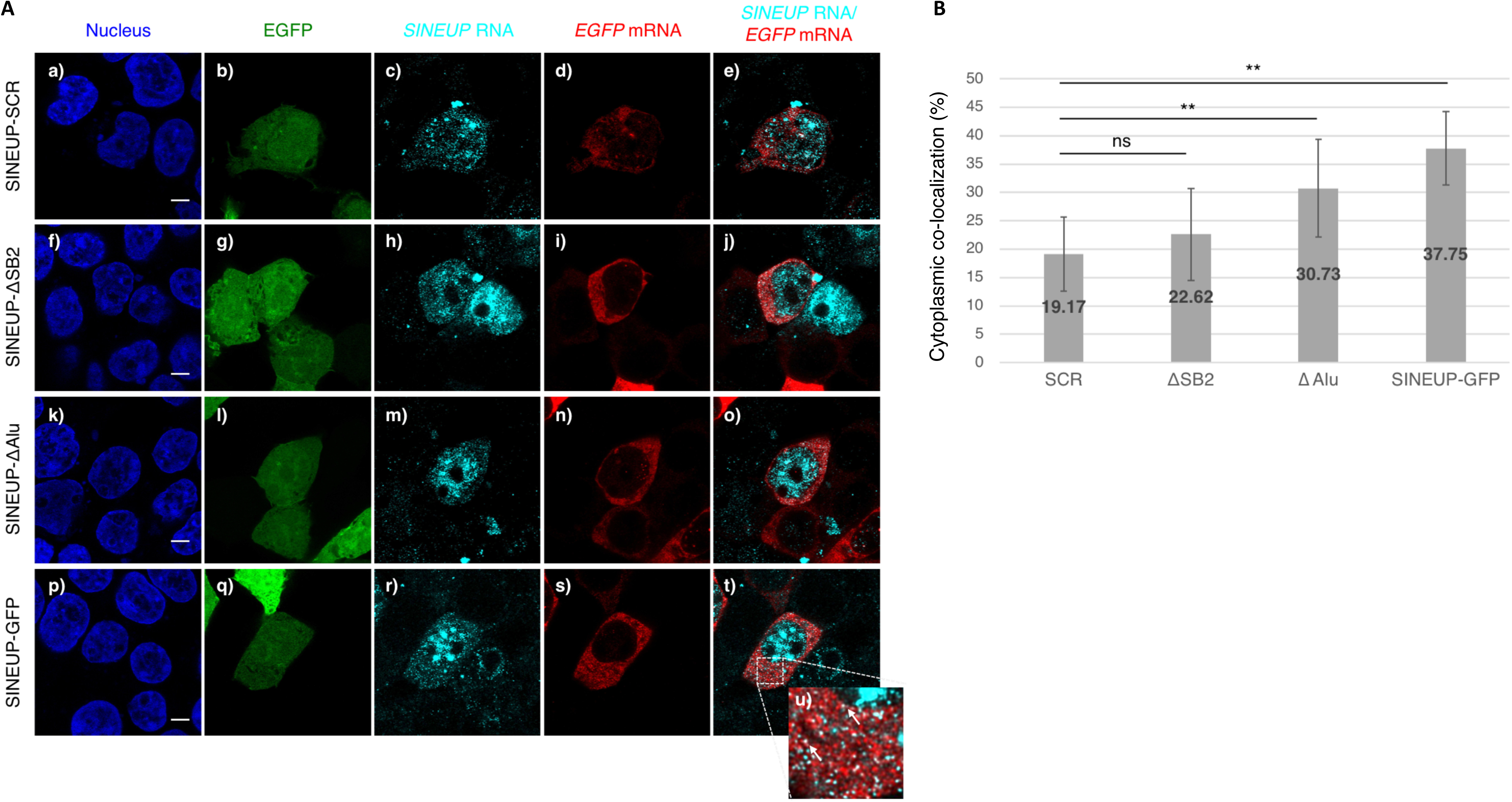
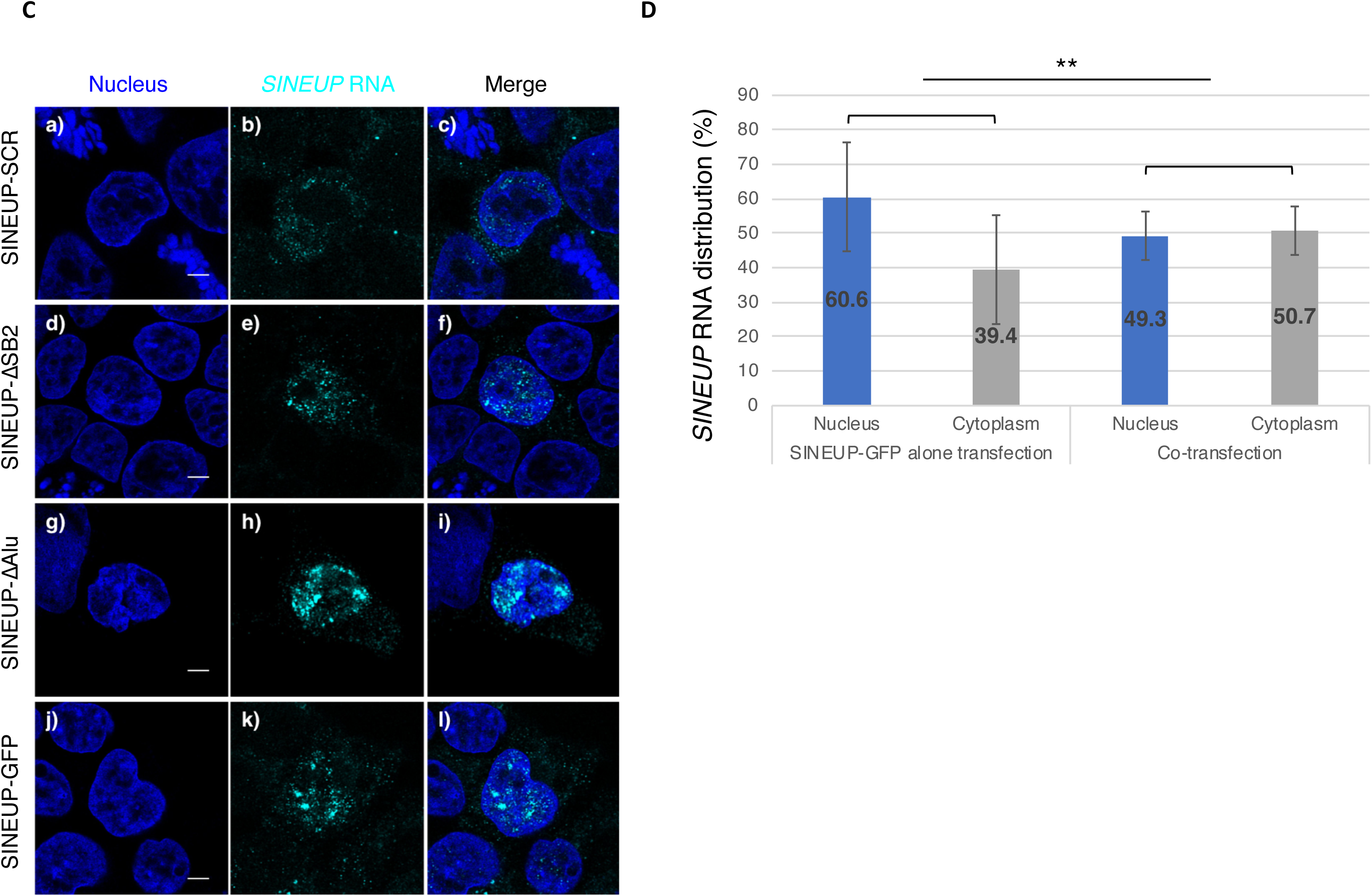
Co-localization of *SINEUP-GFP* RNAs with *EGFP* mRNAs in the cytoplasm. A Subcellular localization of *SINEUP* RNAs and *EGFP* mRNAs. Bars indicate 5 μm. B Comparison of percentage co-localization of *EGFP* mRNAs and *SINEUP* RNAs in the cytoplasm. Data are means ± SD of at least 10 independent cell images. **p < 0.01, ns: not significant by Student’s *t*-test. C Subcellular distribution of *SINEUP* RNAs following transfection with SINEUP expression vectors alone. Bars indicate 5 μm. D Quantitative comparison of the distribution of *SINEUP-GFP* RNA in the presence and absence of *EGFP* mRNA. The ratios of detected spots in the nucleus and cytoplasm were compared between co-transfection of SINEUP-GFP and EGFP vectors, and transfection of SINEUP-GFP vector alone. Data are means ± SD of at least 10 independent cell images. **p < 0.01, ns: not significant by Student’s *t*-test.

#### Identification and functional analysis of *SINEUP* RNA binding proteins

We hypothesized that *SINEUP* RNA binding proteins (RBPs) may play a crucial role in EGFP expression. To identify SINEUP RBPs, we used a modified version of the Chromatin Isolation by RNA Purification (ChIRP) method (Chu *et al*, 2015) (Fig EV4A, B) followed by mass spectrometry (MS) analysis. By carrying out three or more independent experiments on SINEUP-SCR, SINEUP-ΔSB2, SINEUP-ΔAlu and SINEUP-GFP transfection, we detected several SINEUP RBPs. To determine which RBPs are most important for the translational up-regulation activity of SINEUP-GFP, we selected several candidate SINEUP-GFP RBPs with high reliability and specificity (Fig 3) while non-specific bound proteins, which were also detected with beads and LacZ samples, were cut off. We then compared the SINEUP-GFP RBPs with those for the other SINEUP mutants (Fig EV4C-E), and assumed that several nucleocytoplasmic shuttling-related proteins were specifically enriched as SINEUP-GFP RBPs (Fig 3). After excluding ribosomal proteins, we performed siRNA-mediated knockdown of enriched proteins in the scatterplot (Fig 3) to assess their effects on protein up-regulation activity of SINEUP-GFP (Fig 4; Fig EV5A–F). The experiments revealed that knockdown of either PTBP1 (Fig 4A1 and A2) or HNRNPK (Fig 4B1 and B2) significantly decreased the translational up-regulation activity of SINEUP-GFP compared with SINEUP-GFP transfected with scramble siRNA as negative control (Fig 4C1 and C2, SINEUP-GFP ※). Knockdown of EF1A1 decreased the translational-up regulation for EGFP but also affected for β-actin as a non-specific effect (Fig EV5C), suggesting it affected global translational pathways. Interestingly, knockdown of PTBP1 (Fig 4Da-e) and HNRNPK (Fig 4Df-j) significantly reduced the co-localization of *EGFP* mRNA and *SINEUP-GFP* RNA in the cytoplasm (Fig 4E). Furthermore, it also significantly increased *SINEUP-GFP* RNA nuclear retention: 76.4% for PTBP1 knockdown and 61.4% for HNRNPK knockdown versus 49.3% in control cells (Fig 4F) suggesting a decrease in cytoplasmic SINEUP RNA; a smaller portion of cytoplasmic SINEUP RNA co-localized with *EGFP* mRNA. These results suggest that PTBP1 and HNRNPK may participate in the nucleocytoplasmic shuttling of *SINEUP* RNAs.

**Figure 3.**
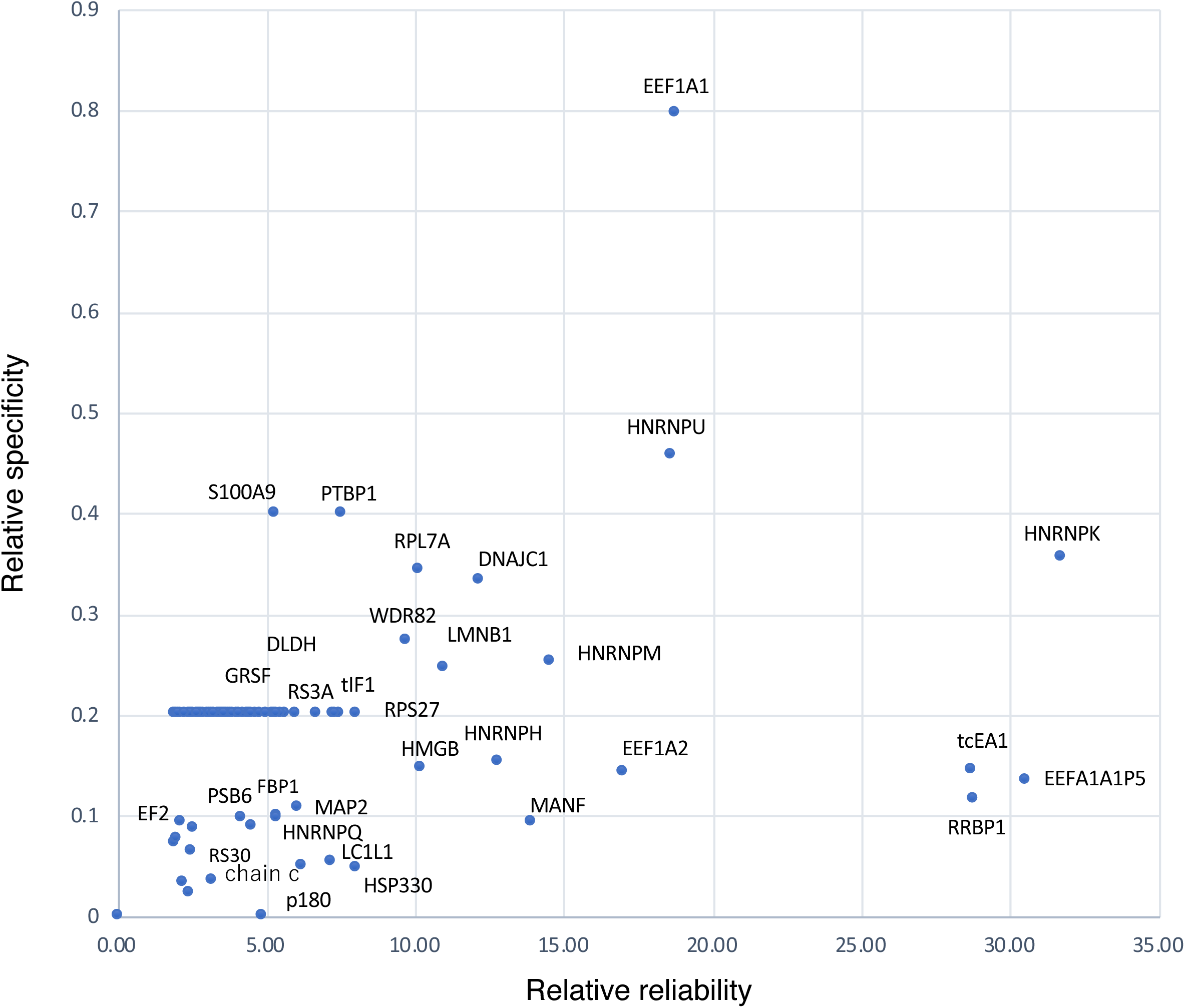
SINEUP-GFP RNA binding proteins. Detected RNA binding proteins (RBPs) were plotted according to relative reliability 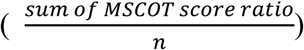 on the x axis, and relative specificity 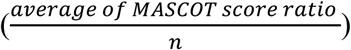 on the y axis. Proteins that were detected by beads and LacZ probes were omitted.

**Figure 4.**
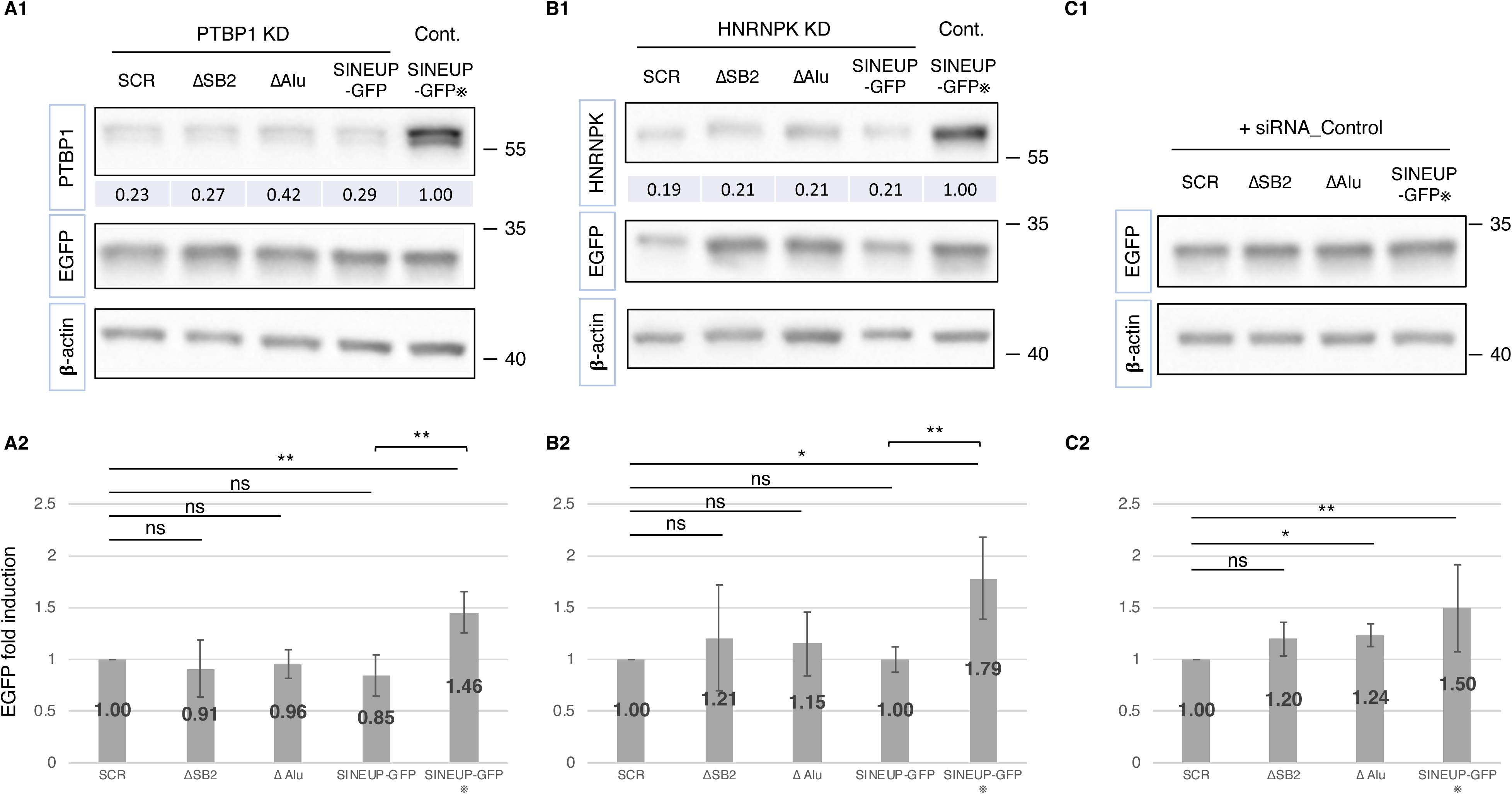
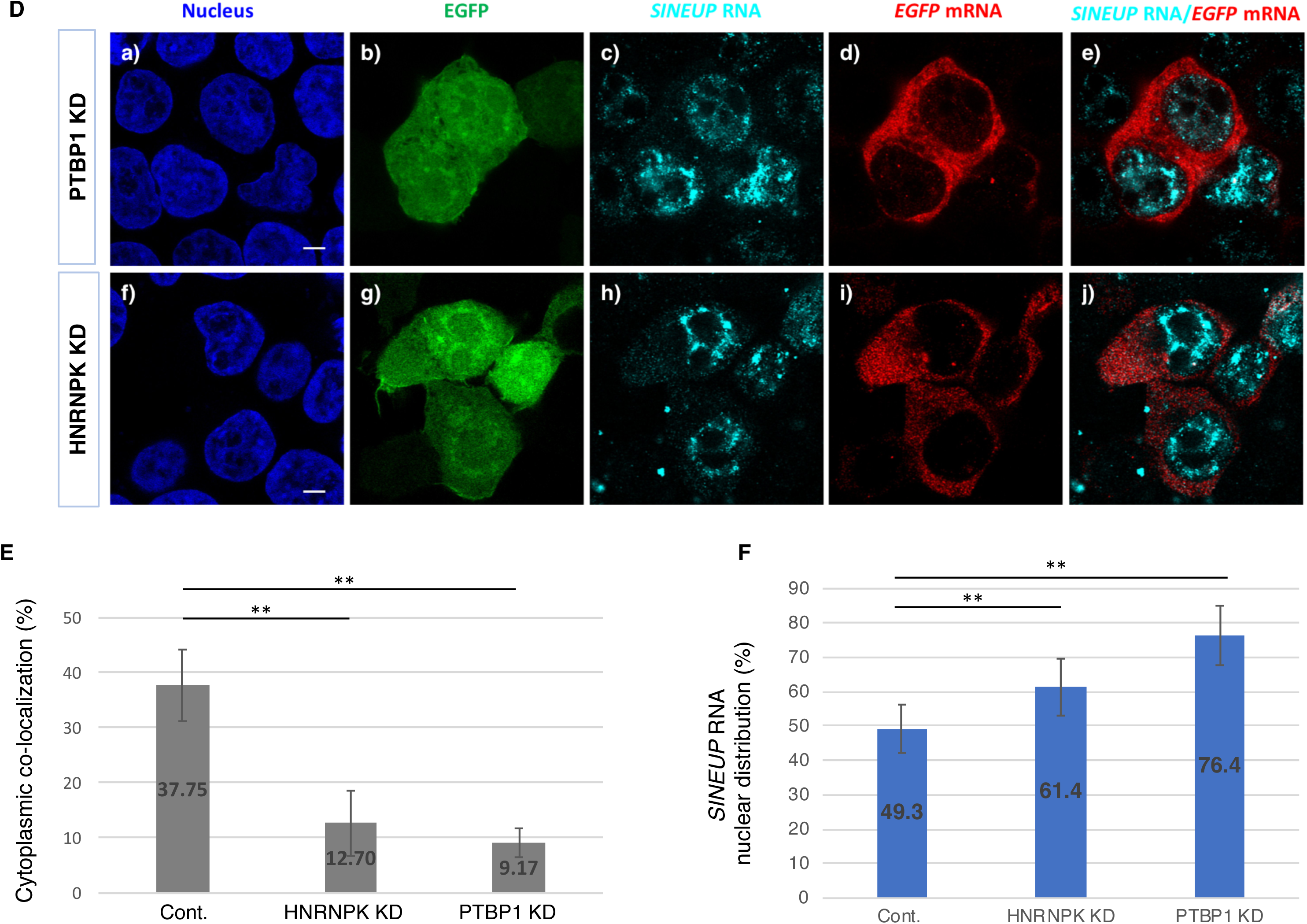
Knockdown of SINEUP RBPs. A1, B1 Representative Western blot images of knockdown (KD) of PTBP1 (A1) and HNRNPK (B1) mediated by siRNA-PTBP1 and siRNA-HNRNPK, respectively. Numbers at the top row under the image show knockdown efficiency compared with cells co-transfected with SINEUP-GFP and negative control siRNA (SINEUP-GFP※ in C1 and C2). C1 Representative Western blot images of transfection with negative control siRNA. A2, B2, C2 Protein levels of EGFP were quantified by Western blot analysis. EGFP expression levels were normalized to that of β-actin. Fold induction of EGFP is calculated relative to cells transfected with the siRNA indicated above panel A1, B1, or C1 respectively. **p < 0.01, *p < 0.05; ns: not significant by Student’s *t*-test. Data are means ± SD of at least 3 independent experiments. D Representative FISH images following knockdown (KD) of PTBP1 (a-e) or HNRNPK (f-j) by siRNAs. Bars indicate 5 μm. E Quantitative comparison of co-localization of *EGFP* mRNAs and *SINEUP* RNAs in the cytoplasm when PTBP1 (D, e) or HNRNPK (D, j) were knocked down. **p < 0.01 by Student’s *t*-test. Data are means ± SD of 10 individual cell images. F Quantitative nuclear distribution of *SINEUP-GFP* RNAs following knockdown of PTBP1 (D, c) or HNRNPK (D, h) by siRNAs; the results are compared with the negative control (without siRNA). For both PTBP1 and HNRNPK, the ratio of *SINEUP-GFP* RNA levels in the nucleus and the cytoplasm were compared between the knockdowns and the negative control. **p < 0.01 by Student’s *t*-test. Data are means ± SD of at least 10 independent cell images.

To better understand the role of PTBP1 and HNRNPK interactions in *SINEUP-GFP* RNA nucleocytoplasmic shuttling, we conducted an RNA immunoprecipitation (RIP) assay of RNA– protein interactions with PTBP1 and HNRNPK proteins in the nucleus and cytoplasm. *SINEUP-GFP* RNAs were pulled down with PTBP1 both in the nucleus (Fig 5A1) and cytoplasm (Fig 5A2) and *EGFP* mRNAs were pulled down with HNRNPK both in the nucleus (Fig 5B1) and cytoplasm (Fig 5B2). These observations suggest that (a) PTBP1 protein was able to bind to *SINEUP-GFP* RNA or the *EGFP-SINEUP* RNA complex in either the nucleus or the cytoplasm, but not to *EGFP* mRNA alone; and (b) HNRNPK protein was able to bind to *EGFP* mRNA or the *EGFP–SINEUP* RNA complex in either the nucleus or the cytoplasm, but not did bind to *SINEUP-GFP* RNA alone. Taken together, these findings indicate that PTBP1 and HNRNPK play a role in the formation of the RNA-protein complexes and participate in the kinetic distribution of these RNA-protein complexes.

**Figure 5.**
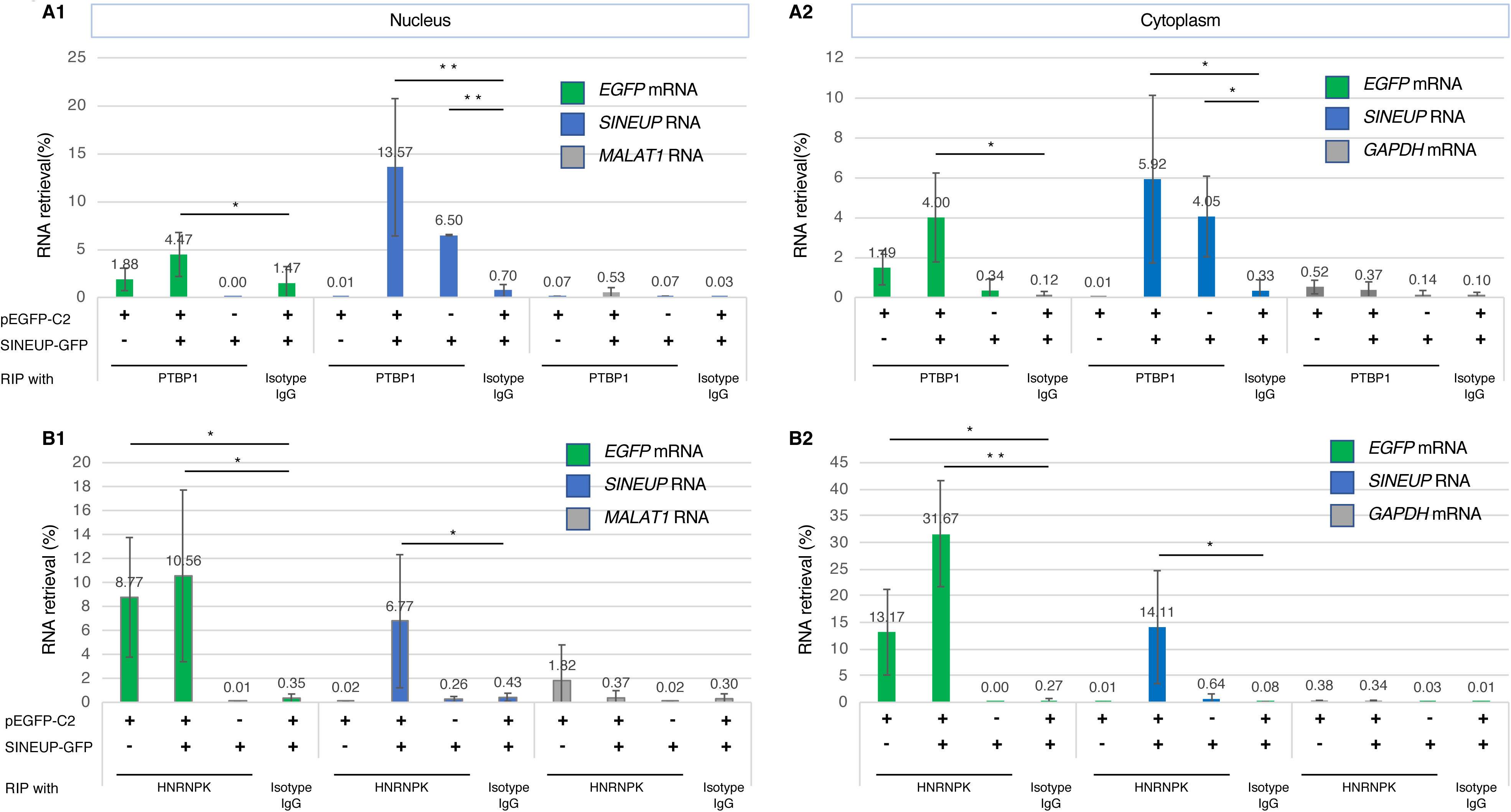
RNA immunoprecipitation. A1, A2 RIP with PTBP1 antibody in the nucleus (A1) and cytoplasm (A2). Isotype IgG was utilized as the negative antibody control. As negative target RNAs, transcripts from the housekeeping genes, *MALAT1* and *GAPDH* were utilized as controls in the nucleus and cytoplasm, respectively. *p < 0.05, **p < 0.01 by Student’s *t*-test. Data are means ± SD of at least 3 independent experiments. B1, B2 RNA immunoprecipitation (RIP) with HNRNPK antibody or isotype immunoglobulin G (Isotype IgG; negative control) in the nucleus (B1) and cytoplasm (B2). The cell lysate co-transfected pEGFP-C2 and SINEUP-GFP vectors or that transfected ether vector alone were tested.

### SINEUP RBPs drive subcellular localization of *SINEUP* RNAs and participate in translational initiation assembly

We next wondered whether EGFP levels would be further enhanced by overexpression of PTBP1 and HNRNPK proteins. To address this question, we transfected the cells with either PTBP1 or HNRNPK expression vector. The EGFP level was moderately but significantly increased by overexpression of PTBP1 (Fig 6A1 and A3) or HNRNPK (Fig 6B1 and B3) in cells co-transfected with EGFP and SINEUP-GFP vectors, but not in those transfected with EGFP vector alone (Fig 6A2, B2). This finding suggests that PTBP1 and HNRNPK formed a protein–*SINEUP* RNA complex and functionally enhanced EGFP levels.

**Figure 6.**
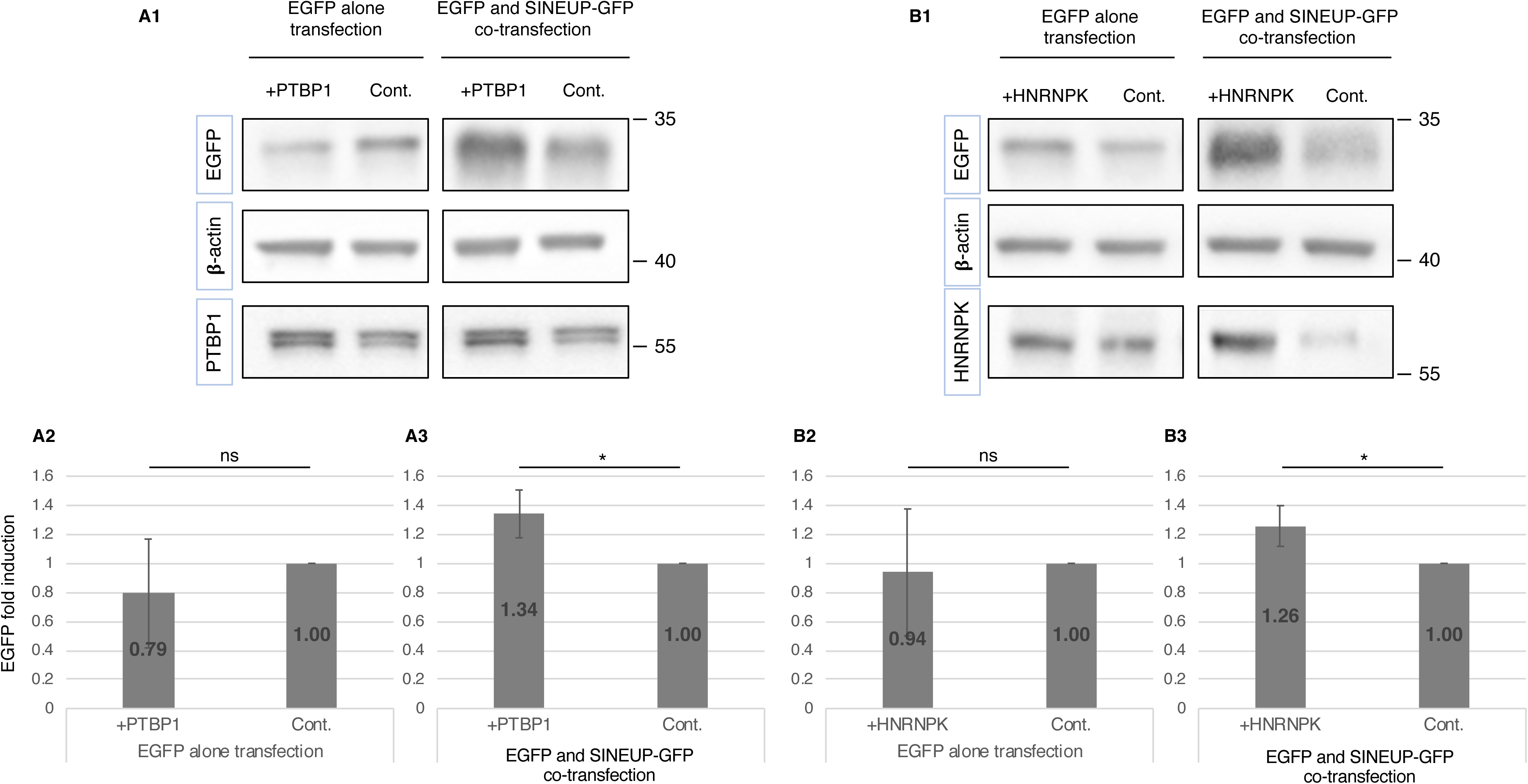
Overexpression of SINEUP RBPs. A1, B1 Representative Western blot images comparing EGFP expression after overexpression of PTBP1 (+PTBP1) (A1) or HNRNPK (+HNRNPK) (B1) with that in the non-overexpressing control (Cont.). A2, A3, B2, B3 Quantification of EGFP levels after non-/overexpression of PTBP1 (A2, A3) or HNRNPK (B2, B3) when cells were transfected with EGFP vector alone (A2, B2) or co-transfected with EGFP and SINEUP-GFP vectors (A3, B3). *p < 0.05, ns: not significant by Student’s *t*-test. Data are means ± SD of at least 3 independent experiments.

To evaluate the effect of SINEUP RBPs on the subcellular distribution of *SINEUP* RNA, we overexpressed (or not) PTBP1 (Fig 7A) or HNRNPK (Fig 7B) in the cells and then co-transfected the cells with EGFP and SINEUP-GFP vectors or with SINEUP-GFP vector alone. Some portion of nuclear *SINEUP-GFP* RNA was shuttled into the cytoplasm when *EGFP* and *SINEUP-GFP* transcripts were co-transfected into cells overexpressing PTBP1 than into cells with normal PTBP1 levels (Fig 7A2); this difference was not seen when the cells were transfected with SINEUP-GFP vector alone (Fig 7A3). Induction of PTBP1 did not directly drive *SINEUP-GFP* RNA from the nucleus to the cytoplasm without the presence of *EGFP* mRNAs. In contrast to PTBP1, overexpression of HNRNPK had no significant effect on the subcellular distribution of *SINEUP-GFP* RNAs (Fig 7B2 and B3). Taking these results together, these findings indicate that PTBP1 and HNRNPK participate in nucleocytoplasmic shuttling of RNA-protein complex and play another role after shuttling into the cytoplasm to regulate translation.

**Figure 7.**
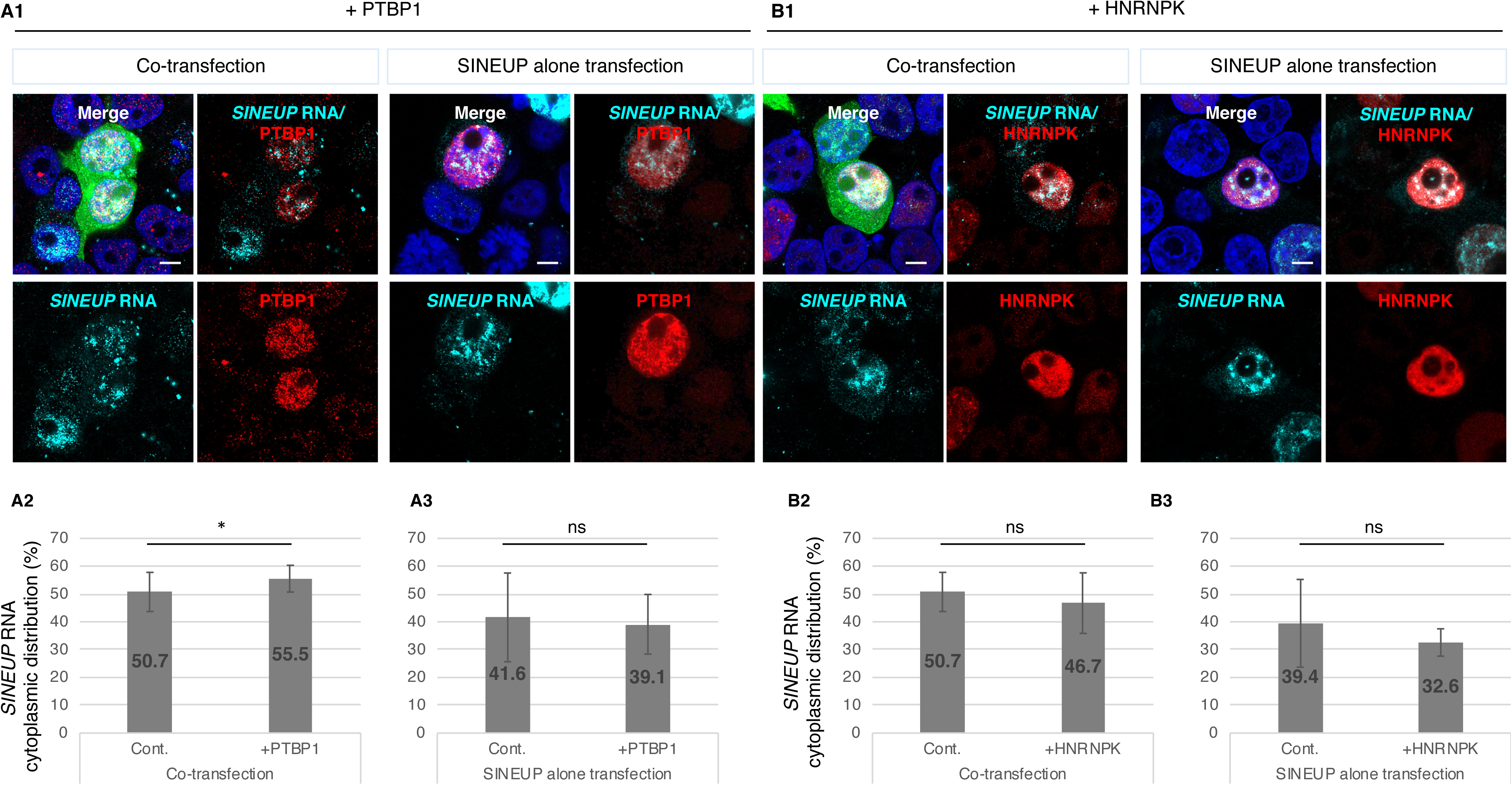
Subcellular distribution of *SINEUP* RNAs after overexpression of SINEUP RBPs. A1, B1 Representative RNA FISH with immunofluorescence images of the subcellular distribution of *SINEUP* RNAs and SINEUP RBPs in cells overexpressing PTBP1 (+PTBP1) (A1) or HNRNPK (+HNRNPK) (B1). Images for cells co-transfected with EGFP and SINEUP-GFP vectors (left images, A1 and B1) were compared with cells transfected with SINEUP-GFP vector alone (right images, A1 and B1). Bars indicate 5 μm. A2, A3, B2, B3 Quantitative comparison of *SINEUP-GFP* RNA distribution between cells overexpressing PTBP1 (A2, A3) or HNRNPK (B2, B3) and non-overexpressing these proteins (Cont.). Results for cells co-transfected with EGFP and SINEUP-GFP vectors (A2 and B2) and those transfected with SINEUP-GFP vector alone (A3, B3) are shown. *SINEUP* RNA signals were detected using Icy Spot Detector. The ratio of spots in the nucleus and the cytoplasm were compared between overexpression and non-overexpression of SINEUP RBPs. *p < 0.05 by Student’s *t*-test. Data are means ± SD of at least 10 independent cell images.

To gain further insights into the molecular mechanism of translational enhancement, we analyzed the distributions of the RNAs and RBPs in polysome fractions of cells overexpressing PTBP1 and HNRNPK. The cytoplasmic lysate was separated into 12 fractions by using a 15%–45% sucrose gradient (Fig 8A). *EGFP* mRNAs were slightly shifted into heavier polysomes when PTBP1 (Fig 8Ba) or HNRNPK (Fig 8Bb) was overexpressed compared with when these SINEUP RBPs were not overexpressed, which is in cells co-transfected with EGFP and SINEUP-GFP vectors (Cont., Fig 8Bc) and in those transfected with EGFP vector alone (Fig 8Bd). *SINEUP-GFP* RNA co-sedimented with *EGFP* mRNA in the heavy polysome fractions when *EGFP* mRNA was present (Fig 8Be, f, g). Although we observed in the FISH analysis (Fig 2C and D) that most *SINEUP-GFP* RNA was retained in the nucleus when the cells were transfected with SINEUP-GFP alone, more than 85% of the cytoplasmic *SINEUP-GFP* RNA sedimented in the fractions containing free/40S binding RNAs (47.7%) to monosomes (37.8%) (Fig 8Bh). This implies that the cytoplasmic *SINEUP-GFP* RNA may participate in an initial phase of translation. We then conducted Western blot analysis of the co-distribution of SINEUP RBPs and the RNAs in the polysome fractions (Fig 8Bi–m). The analysis revealed that HNRNPK co-sedimented with the RNAs and PTBP1 in the light polysome fractions when HNRNPK was overexpressed (Fig 8Bj). PTBP1 also co-sedimented with the RNAs in the light polysome fractions when PTBP1 was overexpressed (Fig 8Bi). Additionally, PTBP1 co-sedimented with *SINEUP-GFP* RNA from the free/40S fractions to the monosome fractions even when SINEUP-GFP vectors were transfected alone (Fig 8Bm). This indicates that *SINEUP* RNAs associate with PTBP1 and may recruit ribosome subunits to contribute to the formation of translational initiation complexes, including elongation factor EF1A, to participate in the initial phases of translation.

**Figure 8.**
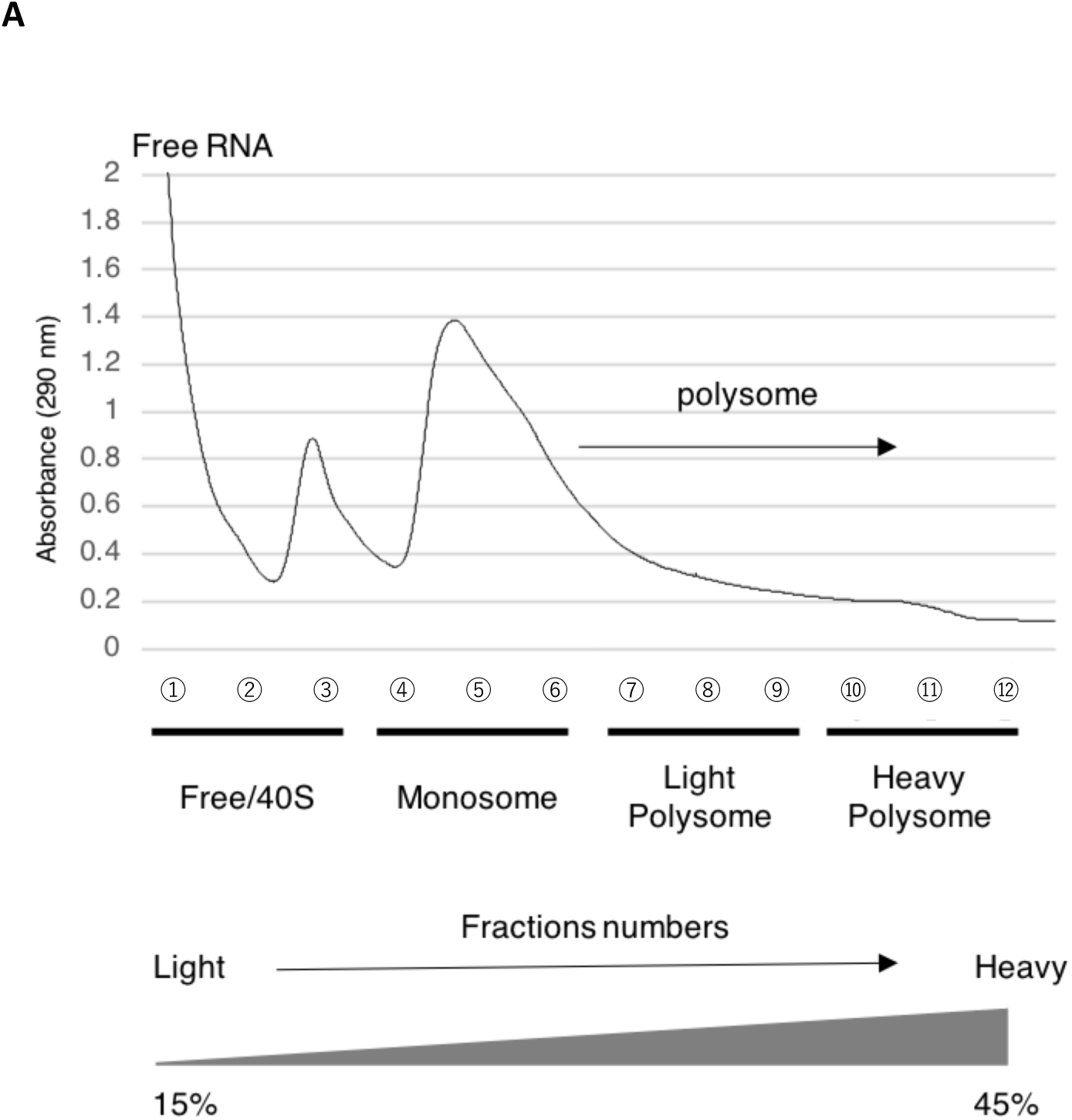
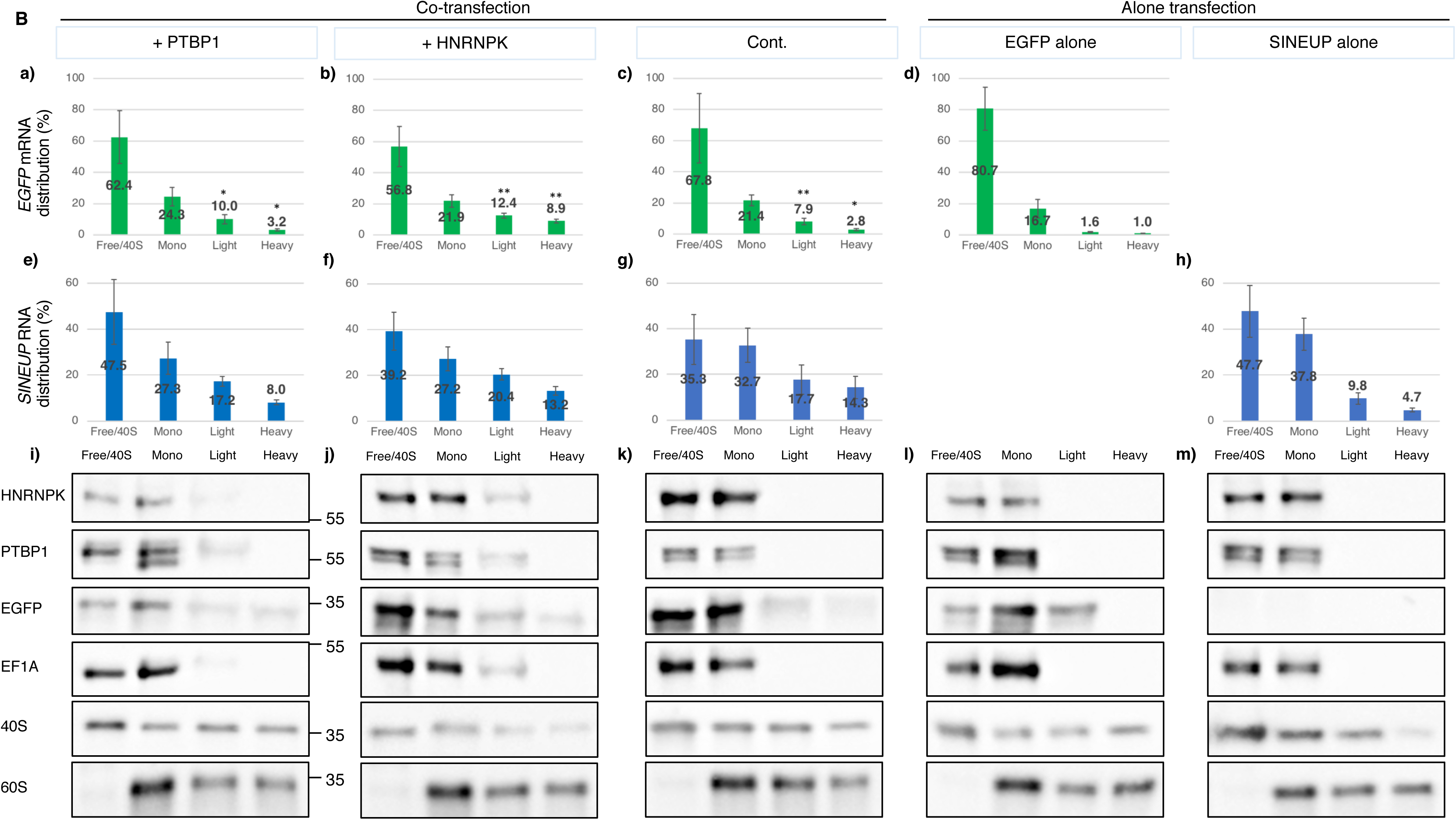
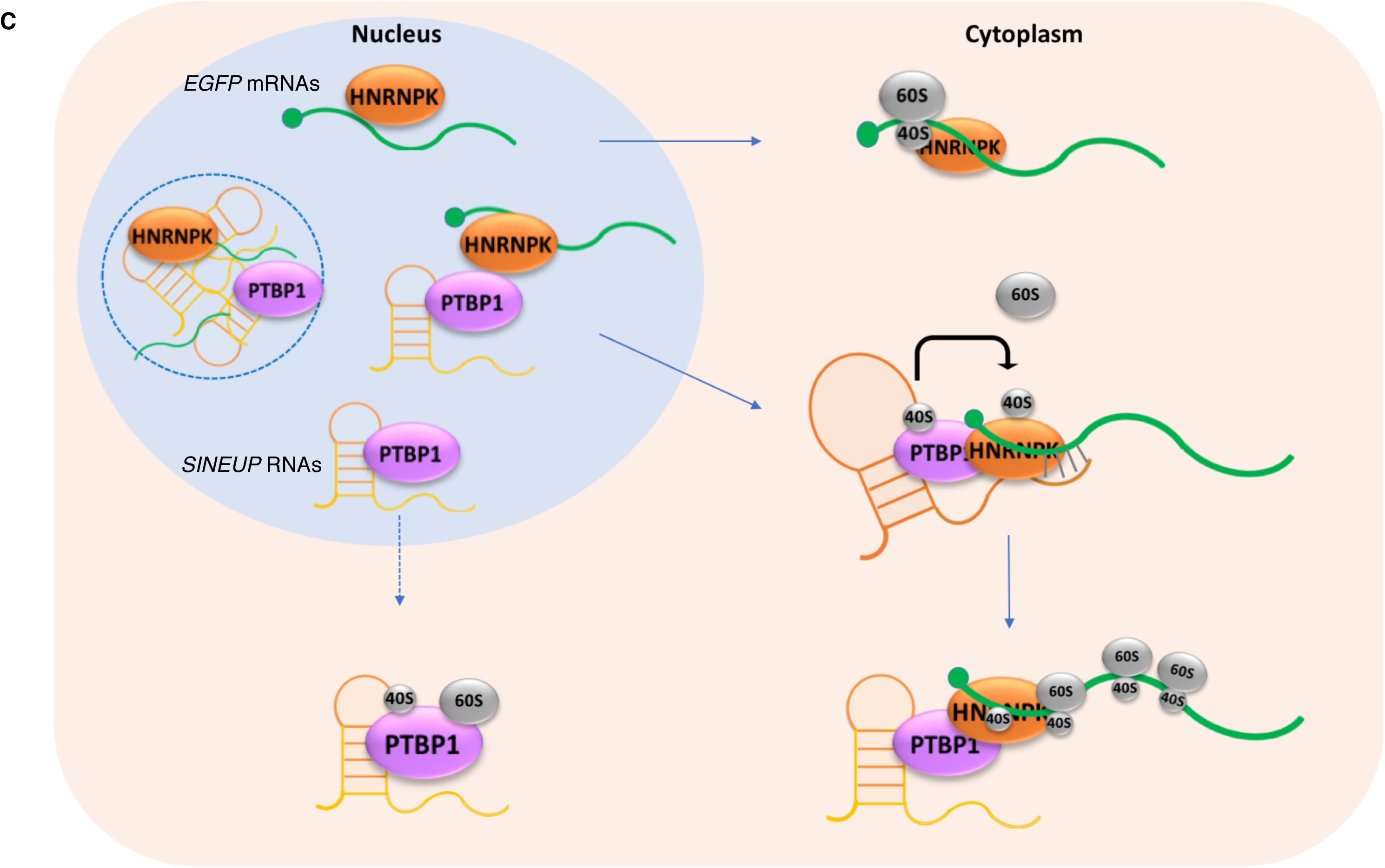
RNA distribution in polysome fractions obtained from cells overexpressing SINEUP RBPs. A Representative polysome gradient profile with optical density (290 nm). Fractions 2460 and ⑫ correspond to the 15% and 45% sucrose fractions, respectively. B Polysome profiling of *EGFP* mRNA (green) and *SINEUP-GFP* RNA (blue). RNA distribution was quantified by RT-qPCR. Each fraction of *EGFP* mRNA in cells co-transfected with EGFP and SINEUP-GFP vectors (a,b,c) was compared with the corresponding fraction in cells transfected with EGFP vector alone (d). *p < 0.05, **p < 0.01 by Student’s *t*-test. Data are means ± SD from 3 independent experiments. The protein distribution in polysome fractions with non-/overexpressed PTBP1 or HNRNPK were tested (i-m). Equal volumes of solution were pooled as Free/40S (Free or 40S binding RNA fraction from ①–③ at A), Mono (monosome fraction from ④–⑥), Light (light polysome fraction from ⑦–⑨), and Heavy (Heavy polysome fraction from ⑩–⑫) and were applied to 10% SDS gels to detect each target protein. Western blot images representative of least 3 independent experiments are shown. C Model of *SINEUP* RNA and SINEUP RBP interactions. SINEUP RBPs (PTBP1 and HNRNPK) participate in *SINEUP* RNA localization both in the nucleus and the cytoplasm. In the nucleus, some portion of the RNAs form RNA–protein granules (dotted circle) where probably immature transcripts accumulate to cluster non-proper conformation, and the other portion of mature transcripts including *EGFP* mRNA and *SINEUP* RNA form complex with SINEUP RPBs, which may be shuttled into the cytoplasm. In the cytoplasm, *SINEUP* RNAs co-operate with the SINEUP RBPs likely re-modelling *SINEUP* RNA structure, and recruit ribosomal subunits to efficiently supply them to *EGFP* mRNA, resulting into positive translational enhancement of *EGFP* mRNA. *EGFP* mRNA can be exported into the cytoplasm by itself, but its translation is initiated more efficiently when *SINEUP-GFP* RNA is present in the cytoplasm.

To investigate direct protein–protein interactions, we performed BS3 chemical cross-linked immunoprecipitation. The results revealed that PTBP1 and HNRNPK directly interacted with each other both in the nucleus and the cytoplasm (Fig EV6A, B), and PTBP1 and HNRNPK also had direct interactions with 40S and 60S ribosome subunits in the cytoplasm, suggesting that these SINEUP RBPs form a complex with *SINEUP* RNAs and *EGFP* mRNAs, and participate together in nucleocytoplasmic shuttling and translational regulation.

## Discussion

In this study, we demonstrated that a synthetic *SINEUP* RNA plays a critical role in the enhancement of the translation of its target mRNA by 1) co-localizing with the target mRNA in the cytoplasm, 2) interacting with SINEUP RBPs to influence the *SINEUP* RNA’s distribution and 3) increasing target mRNA shifting to polysome by participating in translational initiation assembly.

By conducting RNA FISH, we revealed that the co-localization of *EGFP* mRNA and *SINEUP-GFP* RNA in the cytoplasm was required for positive translational regulation. We previously reported that the AS-*Uchl1* is enriched in the nucleus and exported to the cytoplasm under rapamycin-induced stress, which inhibits the mTOR (mammalian target of rapamycin) pathway, resulting in up-regulation of UCHL1 translation (Carrieri et al, 2012); however, in our subsequent study, the distribution of synthetic *SINEUP* RNAs in the cytoplasm was not increased by inhibition of the mTOR pathway (Zucchelli *et al*, 2015). Here, we demonstrated that the synthetic *SINEUP-GFP* RNA localized both in the nucleus and the cytoplasm when *EGFP* mRNA and *SINEUP* RNA co-existed, but was retained in the nucleus in the absence of *EGFP* mRNA. This suggests that synthetic *SINEUP* RNAs may have specific export systems that rely on specific unknown motifs for SINEUP-binding factors, and which differ from the mTOR-dependent export system of AS-*Uchl1*. From loss- and gain-of-function studies of SINEUP RBPs, we found that PTBP1 and HNRNPK proteins have key roles in *SINEUP-GFP* RNA nucleocytoplasmic shuttling and in up-regulating translation.

PTBP1 (also known as HNRNPI) is a multifunctional RNA binding protein that participates in alternative splicing, mRNA stabilization, and nucleocytoplasmic shuttling by binding to polypyrimidine-rich tract in pre-mRNAs (Singh *et al*, 1995; Sawicka *et al*, 2008; Romanelli *et al*, 2013). PTBP1 is known as a binding factor for TOP mRNA, which contains 5’ terminal oligopyrimidine tract (5’TOP) mostly found in mRNAs encoding ribosomal protein and elongation factors (Avni *et al*, 1997; Meyuhas, 2000), to regulate translation of the target TOP mRNA (Gismondi *et al*, 2014) as a *cis*-acting regulator. LARP1, which is known as 5’TOP mRNA binding proteins, stabilize the target 5’TOP mRNAs by forming complex with 40S ribosome subunit (Gentilella *et al*, 2017), In this study, SINEUP-GFP showed direct interaction with 40S, implying that SINEUP RNA with multimerization including PTBP1, 40S and the target mRNA may contribute the target mRNA stabilization. There are several reports that PTBP is recruited to cap-independent translation to stimulate translational initiation with re-modelling the target transcripts followed by supplying the small subunit (40S) to binding sites (Caceres *et al*, 2016; Stoneley & Willis, 2004). In one example of translational initiation mechanism, the cricket paralysis virus (CrPV) recruits 40S ribosome subunit on its RNA with high affinity, and requires eukaryotic elongation factor (eEF)1A and eEF2 to recruit tRNA for assembly of 80S tertiary complexes (Johnson *et al*, 2017). One of the key findings in this study, several components of ribosomal complexes such as eEF1A, eEF2 and ribosome proteins were also detected as SINEUP RBPs (Fig 3), and EF1A and PTBP1 were co-sedimented with the RNAs in various fractions ranging from free/40S-binding RNA to light polysome fractions. This indicates that *SINEUP-GFP* RNA may become a scaffold to bind PTBP1, and to recruit translational initiation factors; this likely supplies the 40S subunit to mRNA binding sites, thereby helping initiation assemblies (Fig 8C). Although we need to further characterize the protein–protein interactions, our data suggest that complexes of SINEUP RBPs plus *SINEUP* RNAs affect translational regulation in the initiation phases.

HNRNPK has three KH (K homology) domains, which bind RNAs, and unique nuclear localization signals which have bi-directional transport, that enable to export the nuclear envelope with target mRNAs (Michael *et al*, 1997). HNRNPK regulates the target mRNA’s translation positively or negatively, depending on the target mRNA. As an example of positive regulation, HNRNPK bound to *VEGF* mRNA and stimulated the ribosome to bind the mRNA resulting in a shift to heavier polysomes (Feliers *et al*, 2007). In contrast, in a case of negative regulation, HNRNPK blocked monosome assembly by binding to the 3’ UTR of *c-Src* mRNA thereby repressing the translation (Naarmann *et al*, 2008). In our study, HNRNPK shifted to heavier polysomes with *EGFP* mRNA when HNRNPK was overexpressed and up-regulated *EGFP* mRNA translation, suggesting positive regulation; HNRNPK contributed to ribosome assembly only when *EGFP* mRNA and *SINEUP-GFP* RNA co-existed.

When PTBP1 or HNRNPK were overexpressed, EGFP enhancement was observed only when *SINEUP-GFP* RNA and *EGFP* mRNA coexisted, i.e., not when the cells were transfected with *EGFP* construct alone. Similarly, when PTBP1 was overexpressed, *SINEUP-GFP* RNA distribution was affected only when *SINEUP-GFP* RNA and *EGFP* mRNAs co-existed. These findings imply that the SINEUP RBPs, *SINEUP-GFP* RNA, and *EGFP* mRNAs together form RNA–protein complexes, and that these multiple components are essential for a functional complex. Several studies report protein–protein direct interaction between PTBP1 and HNRNPK, suggesting that they function cooperatively in biological processes (Kim *et al*, 2000). One lncRNA, *TUNA*, which is known to maintain pluripotency in neuronal cells, to form complexes including PTBP1 and HNRNPK, associates with *Nanog, Sox2*, and *Fgf4* to activate these pluripotency genes (Lin *et al*, 2014). *Lncenc1*, a highly abundant lncRNA in naive embryonic stem cells, recruits PTBP1 and HNRNPK and to bind to glycolysis genes, thereby regulating cell pluripotency and glycogenesis (Sun *et al*, 2018). Taken together, our results and these reports indicate that formation of RNA– protein complexes containing *SINEUP-GFP* RNAs and RBPs such as PTBP1 and HNRNPK affects not only nucleocytoplasmic shuttling of the *SINEUP-GFP* RNAs, but also participates in translational initiation assembly to recruit initiation factors for up-regulating translation of the target mRNA (Fig 8C).

Interestingly, both HNRNPK and PTBP1 are classed as heterogeneous nuclear ribonucleoproteins (RNPs), which mainly participate in alternative mRNA splicing, conformation of RNP assembly to compact transcripts in the nucleus, and nucleocytoplasmic shuttling (Dreyfuss *et al*, 1988). RNA–RNA intermolecular interactions in RNP granules, which are non-membrane-bound organelles including specific RBPs, contribute to various cellular functions (Van Treeck & Parker, 2018). Here, we observed that some of the *SINEUP-GFP* RNA was retained in the nucleus and co-localized with the RBPs, suggesting that the *SINEUP* RNA and RBPs may form RNA–protein granules that accrue in the nucleus as part of a yet-unknown mechanism for RNA modification and editing. Some reports suggest that nuclear history determines a transcript’s fate in the cytoplasm (Matsumoto *et al*, 1998), and the exon junction complex, which includes splicing factors such as RNPs, affects mRNA destination (Giorgi & Moore, 2007).

Although the binding domain and structural conformation of binding factors in *SINEUP*– protein interactions require further study, our results suggest that intermolecular interactions between *SINEUP* RNAs and RBPs contribute to the translational up-regulation of the target mRNA. This improved understanding of the mechanisms of efficient protein translational regulation by functional lncRNAs, and will help to facilitate broad applications of RNA regulation such as nucleic-acid–based therapies.

## Materials and Methods

### Cell culture

Human Embryonic Kidney (HEK) 293T/17 cells purchased from ATCC were maintained in DMEM, high glucose, GlutaMAX™ Supplement, pyruvate (Gibco) supplemented with 10% fetal bovine serum (Sigma) and 1% penicillin-streptomycin (Wako) at 37°C, 5% CO_2_.

### Plasmid and constructs

The pEGFP-C2 vector (expression vector for EGFP) was purchased from Clontech. SINEUP-GFP in pcDNA3.1 (-) vector was described previously in Carrieri *et al*, 2012). SINEUP-SCR, SINEUP-ΔSB2, SINEUP-ΔAlu and SINEUP-GFP constructs were cloned into the pCS2+ vector (Indrieri *et al*, 2016).

### Plasmid transfection and conditions

The pEGFP-C2 and SINEUP vectors were co-transfected into HEK293T/17 cells by using Lipofectamine 2000 (Invitrogen) with OptiMEM (1×) Reduced Serum Medium (Gibco). After testing *EGFP* mRNA translation at various time points after transfection, we confirmed that up-regulation of translation of *EGFP* mRNA occurred from 24 h to 48 h post-transfection.

### Measuring protein up-regulation by Western blot assay

Cells were plated in six-well plates, transfected with plasmid(s), lysed in Cell Lysis buffer (Cell Signaling), and incubated at 4°C for 1 h. The cell lysates were applied to 10% precast polyacrylamide gels (Bio-Rad) for SDS-PAGE and transferred to nitrocellulose membranes (Amersham). The membranes were incubated for 1 h at room temperature with the primary antibody, anti-GFP rabbit polyclonal antibody (1:1000 dilution; A6455, Thermo Fisher Scientific), and then for 45 min at room temperature with the secondary antibody, anti-rabbit IgG conjugated with HRP (P0448, Dako), and EGFP was detected by ECL detection reagent (Amersham). As a control, anti-β actin mouse monoclonal antibody (1:1000 dilution; A5441, Sigma Aldrich) was used as the primary antibody, and anti-mouse IgG–conjugated HRP (1:1000 dilution; P0447, Dako) was used as the secondary antibody.

### RNA extraction and reverse transcription real-time quantitative PCR (RT-qPCR)

RNAs were extracted with an RNeasy mini kit (QIAGEN) following the manufacturer’s instructions. The TURBO DNA-free Kit (Invitrogen) was used for DNase I treatment to remove plasmid DNAs. For RT-qPCR, cDNA was synthesized by using the PrimeScript 1st strand cDNA synthesis kit (TAKARA), and PCR was performed with SYBR Premix Ex *Taq* II (TAKARA) and the 7900HT Fast Real-Time PCR System (Applied Biosystems). The thermocycling protocol was 95°C for 30 s followed by 40 cycles of 95°C for 5 s and 60°C for 30 s.

### RNA FISH

FISH probes for target transcripts were designed by using Stellaris RNA FISH designer (BIOSEARCH Technologies; https://www.biosearchtech.com/support/tools/design-software/stellaris-probe-designer), and fluorescently labelled with Quasar 570 (for *SINEUP* RNAs) or Quasar 670 (for *EGFP* mRNA). Cells were fixed with 4% paraformaldehyde (WAKO) and permeabilized with 0.5% Triton X-100 (Sigma) at room temperature for 5 min. Hybridization was performed overnight at 37°C. Nuclei were visualized by incubation with Hoechst 33342 (H3570, Thermo Fisher Scientific). After sequential washing steps, cell images were detected by using a SP8-HyVolution confocal laser scanning microscope (Leica Microsystems) with a 63x/1.4 oil objective lens; the images were processed using HyD detectors with Huygens Essential software (Scientific Volume Imaging). The RNA signals in the images were counted by using Icy Spot Detector (http://icy.bioimageanalysis.org/plugin/Spot_Detector; Olivo-Marin, 2002), and percentage co-localization was calculated by using Icy Colocalization Studio (http://icy.bioimageanalysis.org/plugin/Colocalization_Studio; Lagache *et al*, 2015).

#### Detection of SINEUP RBPs

The protocol for detecting SINEUP RBPs was based on the protocol for the Magna ChIRP RNA Interactome kit (Merck Millipore) with the modification of cross-linking with 300 mJ/cm of 264 nm UV light (CL-1000 Ultraviolet Cross Link, UVP). The cell pellet (2 × 10^7^ cells) was suspended wit 2 mL of Lysis buffer (Cell Signaling) supplemented with Protease Inhibitor Cocktail Set III (Merck Millipore), mixed by rotation at 4°C for 30 min, and then sonicated for 8 cycles (ON for 30 s, OFF for 30 s) using a Picoruptor Sonicator (Diagenode). Each tube of lysate was incubated overnight at 37°C in hybridization buffer (750 mM NaCl, 50 mM Tris-HCl (pH 7.5), 1 mM EDTA) with the addition of 15% v/v formamide (Sigma), Phenylmethanesulfonyl Fluoride (PMSF) (Cell Signaling), Protease Inhibitor Cocktail Set III (Merck Millipore) and SUPERase• In RNase Inhibitor (Thermo Fisher Scientific) just before use, and 100 pmol of probe. Each lysate was then incubated at 37°C for 30 min with washed MagCapture Tamavidin 2-REV magnetic beads (WAKO). After sequential washes, the bead samples were separated into two halves for protein and RNA extraction. Proteins were extracted as reported previously (Chu et. al, 2015); the eluent was incubated in DNase/RNase solution (100 μg/mL RNase A, 0.1 U/μL RNase H, and 100 U/mL DNase I) at 37°C for 1 h followed by acetone precipitation. The protein samples were digested with 10 ng/μL Sequencing Grade Modified Trypsin (V5111, Promega) overnight, and the resultant peptides were subjected to Liquid chromatography–tandem mass spectrometry (LC-MS/MS) at the Support Unit for Bio-Material Analysis, Research Resource Center, Brain Science Institute in Wako, Japan. Proteome Discover (version 1.4) software with the MASCOT search engine (version 2.6.0) was used in the Swiss-Prot database. For RNA extraction, the beads were incubated with proteinase K at 55°C overnight, and then extracted with Trizol (Thermo Fisher Scientific) and chloroform (WAKO). The eluent was treated with DNase I (Ambion), and the RNA expression level was quantified by RT-qPCR.

#### Validation of SINEUP RBPs by siRNA-mediated knockdown

All siRNAs listed below were purchased from Thermo Fisher Scientific. DNAJC1 Silencer Select Pre-designed siRNA (ID: s34557), EEF1A1 Silencer Select Pre-designed siRNA (ID: s4479), EEF2 Silencer Select Pre-designed siRNA (ID: s4492), HNRNPK Silencer Select siRNA: Standard (ID: s6739), HNRNPM Silencer Select Pre-designed siRNA (ID: s9259), HNRNPU Silencer Select Pre-designed siRNA (ID: s6743), LMNB1 Silencer Select Pre-designed siRNA (ID: s8226), and PTBP1 Silencer Select Validated siRNA (siRNA ID: s11434) were utilized for the knockdown experiments; 4390843 Silencer Select Negative Control #1 siRNA was used as the negative control. Twenty-four hours after the cells were plated, the target pre-designed siRNA was transfected by using Lipofectamine RNAiMAX (Invitrogen), and the cells were maintained in DMEM (1×) +GlutaMAX-1 (Gibco) supplemented with 10% fetal bovine serum (Sigma) without penicillin-streptomycin (Wako) at 37°C, 5% CO_2_ for 24 h. Then pEGFP-C2 and/or SINEUPs vectors were transfected into the cells as described above. Targeted proteins were detected by using the following anti-mouse monoclonal antibodies purchased from Santa Cruz: DnaJC1[D-10] (sc-514244), EF-1 α1 [CBP-KK1] (sc-21758), EF-2 [C-9] (sc-166415), hnRNP K/J [3C2] (sc-32307), hnRNP M [A-12] (sc-515008), hnRNP I [SH54] (sc-56701), hnRNP U [3G6] (sc-32315), and Lamin B1 [8D1] (sc-56144). The above primary antibodies were diluted 1 in 500 and then incubated at 4°C overnight. HRP-conjugated anti-mouse IgG (P0447, Dako) was used as secondary antibody diluted 1 in 1000 and then incubated at room temperature for 45 min for protein visualization.

### RIP with SINEUP RBPs

RIP was performed with the Abcam protocol (https://www.abcam.com/epigenetics/rna-immunoprecipitation-rip-protocol) with some modifications. The cells were plated into 10-cm plates, and followed by plasmid transfection described above, and then next day, nuclei and cytoplasmic fractions were isolated. Nuclear pellets were sheared by sonication with 5 cycles (ON for 30 s, OFF for 30 s) using a Bioruptor Pico device. To immunoprecipitate RNA with the antibodies for target proteins, each lysate was incubated at 4°C overnight. Then Protein A/G magnetic beads (Invitrogen) were added to bind the target antibodies, and sequential washing was conducted to remove unbound antibodies. The anti-hnRNP K mouse monoclonal antibody [3C2]-ChIP Grade (ab39975, Abcam) and anti-PTBP1 mouse monoclonal antibody (32-4800, Thermo Fisher Scientific) were used to immunoprecipitate HNRNPK and PTBP1, respectively, in each nuclear or cytoplasmic fraction. To purify the RNA, the lysates were incubated with protease K at 55°C overnight followed by Trizol (Thermo Fisher Scientific) and chloroform (WAKO) extraction. RNA levels were quantified by RT-qPCR.

#### Clone overexpression of SINEUP RBPs

Clone vectors, hnRNPK in pCMV6-XL5, and PTBP1 in pCMV6-AC, were purchased from ORIGENE. After the cells were plated for 18 h, target protein clone vector was transfected by using Lipofectamine 2000 (Invitrogen). The transfected cells were maintained in DMEM (1×) +GlutaMAX-1 (Gibco) supplemented with 10% fetal bovine serum (Sigma) without penicillin-streptomycin (Wako) at 37°C, 5% CO_2_ for 6 h. Then pEGFP-C2 and SINEUP vectors were transfected into the cells as described above.

#### Immunofluorescence microscopy

The cells were prepared as described above for RNA FISH. After the cells were permeabilized, the primary antibodies, anti-hnRNP K mouse monoclonal antibody [3C2]-ChIP Grade (ab39975, Abcam) and anti-PTBP1 mouse monoclonal antibody (32-4800, Thermo Fisher Scientific), were added and hybridized overnight. Alexa Fluor 647–conjugated goat anti-mouse IgG secondary antibodies (A-21236, Thermo Fisher Scientific) were used to visualize the results.

#### Polysome fractionation

Polysome fractionation was performed as reported previously (Faye *et al*, 2014). Briefly, 2.5 × 10^6^ cells were plated into 10-cm plates, 18h later, followed by hnRNPK and PTBP1 clone vector transfection, then additional 6 h later, EGFP and SINEUP vector transfection were performed as described above. The transfected cells were maintained in growth media without antibiotics for 48 h after the clone vector transfection, then incubated with 0.1 mg/mL cycloheximide for 5 min at 37°C followed by washing with ice-cold PBS containing 0.1 mg/mL cycloheximide. The harvested cells were centrifuged at 300 × *g* for 10 min at 4°C. The cell pellets were suspended with 200 μL of ice-cold lysis buffer (50 mM Tris-HCl pH 7.5, 100 mM NaCl, 30 mM MgCl_2_, 0.1 mg/mL cycloheximide, 0.1% NP-40, with fresh RNase inhibitor and Proteinase inhibitor cocktail added just before use). The cell lysate was incubated for 10 min on ice followed by centrifugation at 2,000× *g* at 4°C for 5 min to separate the nuclei. The cytoplasmic fraction was subjected to further centrifugation at 17,000× g at 4°C for 5 min to remove cell debris. The cytoplasmic lysate was layered onto a 15%– 45% sucrose gradient and centrifuged in an SW41Ti Beckman rotor at 190,000× g at 4°C for 3.5 h. The sucrose gradient was separated into 12 fractions calculated by Triax flow cell (Biocomp). Half of each fraction was treated with Proteinase K at 55°C overnight then followed by Trizol and chloroform extractions as described above. The eluent was treated with DNase I (Ambion), and the RNA expression level was quantified by RT-qPCR.

Using the other half of each fraction, proteins were isolated using the Thermo Fisher Scientific acetone precipitation protocol: 4 volumes of cold acetone were added to each sample, and then the samples were incubated at -20°C for 1 h or overnight, and centrifuged at 13,000 × *g* for 10 min at 4°C. Each pellet was suspended in PBS and subjected to Western blot analysis.

#### Protein-protein direct interaction with chemical cross-linking

BS3 (bis [sulfosuccinimidyl] suberate) cross-linking was performed based on a published protocol (de la Parra *et al*, 2018). The cells were incubated with 0.6 mM BS3 (Thermo Fisher Scientific) for 30 min; the reaction was quenched by incubation with 1M Tris-HCl (pH 7.5) for 15 min at room temperature. The cell lysate was incubated with target antibodies as described for RIP. After sequential washes, the beads were incubated with 2× Laemmli sample buffer (Bio-Rad) at 95°C for 20 min to dissociate proteins, followed by Western blot assay.

#### Statistical analysis

Statistical differences were measured using Student’s *t-*test with paired, two-sided. Bar graphs were described as means ± standard deviation (SD) from at least 3 independent experiments. Statistically significances were represented with \**p* < 0.05, \*\**p* < 0.01. To test normality of all data sets, Kolmogorov–Smirnov test was used.

## Acknowledgements

We thank members of PC’s laboratory and of the SINEUP network (SISSA, IIT, University of Eastern Piedmont, RIKEN) for thought-provoking discussions. We thank the Support Unit for Bio-Material Analysis, Research Resource Centre, Brain Science Institute in Wako, Japan for performing Mass spectrometry analysis. This work was partly supported by a Research Grant from the Japanese Ministry of Education, Culture, Sports, Science and Technology (MEXT) to the RIKEN Centre for Integrative Medical sciences and RIKEN Centre for Life Science Technology, by a Basic Science and Platform Technology Program for Innovative Biological Medicine no. 18am0301014, from the Japan Agency for Medical Research and Development (AMED) to PC, and by a JSPS KAKENHI Grant Number JP18K14631 to HT.

## Author contributions

NT performed all experimental work. NT, HT, SZ, SG, and PC planned the work and evaluated the data. NT, HT, and PC wrote the manuscript.

## Conflict of interest

SZ, SG, and PC are inventors on patent US9353370B2 and related applications in EU and JP, and HT, SZ, SG and PC are inventors on patent application IT02018000002368 and related application PCT/IB2019/050914 hold by SISSA and TransSINE Technologies Inc. on SINEUPs technology. PC and SG are founders of TransSINE Technologies Inc., located in Japan. SG and PC are founders of TranSINE Therapeutics, located in UK, and HT and SZ own shares. These conflicts of interests are not competing any commercial interests in relation to this work

## Expanded View Figure legends

**Fig EV1.**
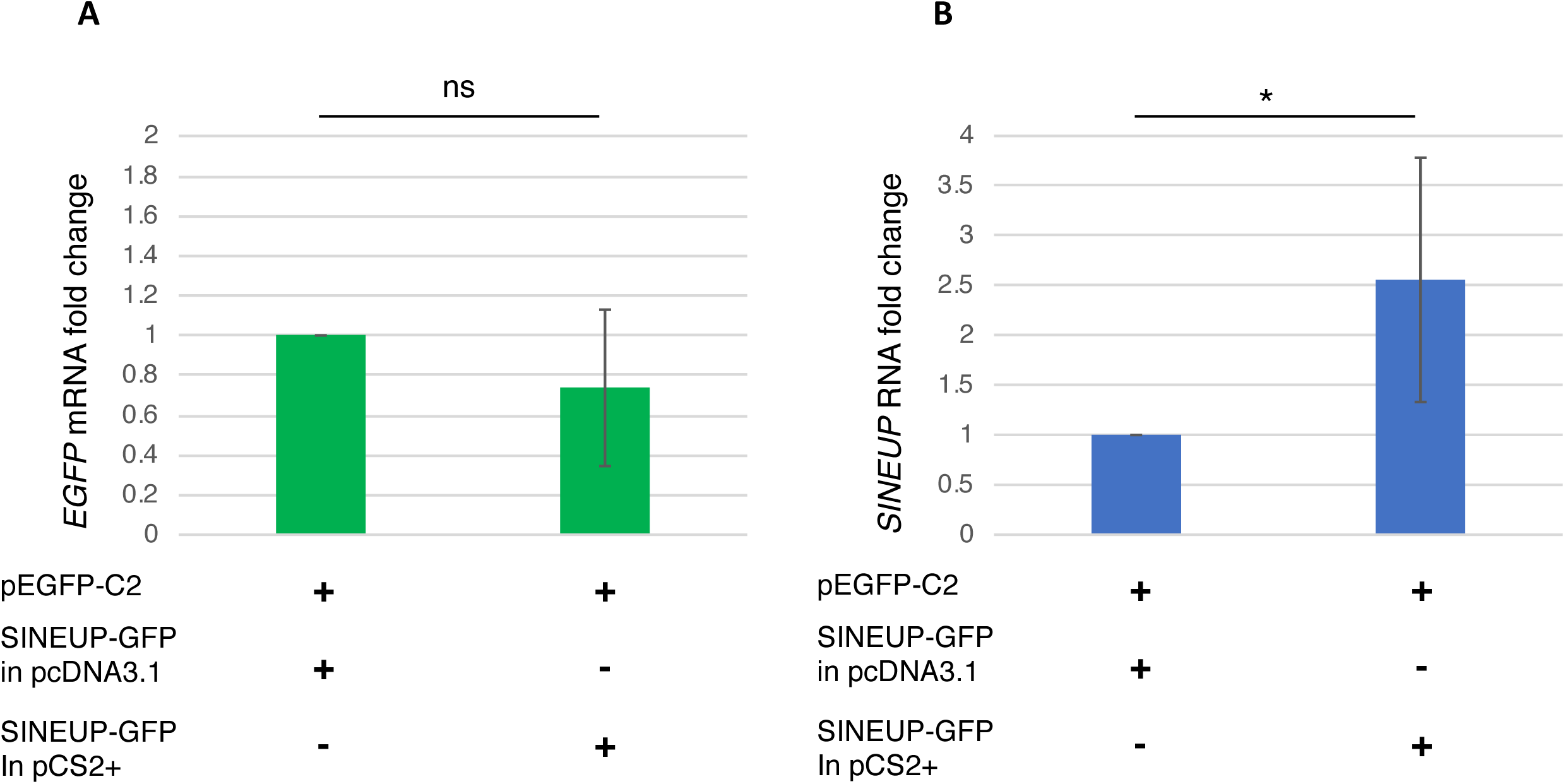
Comparison of the RNA expression level in different plasmid for *SINEUP-GFP*. A Quantitative comparison of *EGFP* mRNA levels in the cells co-transfected EGFP vector and ether SINRUP-GFP in pcDNA3.1 vector or in pCS2+ vector. B Quantitative comparison of *SINEUP-GFP* RNA levels in the cells co-transfected EGFP vector and ether SINRUP-GFP in pcDNA3.1 vector or in pCS2+ vector. *p < 0.05, ns: not significant by Student’s *t*-test. Data are means ± SD of at least 3 independent experiments.

**Fig EV2.**
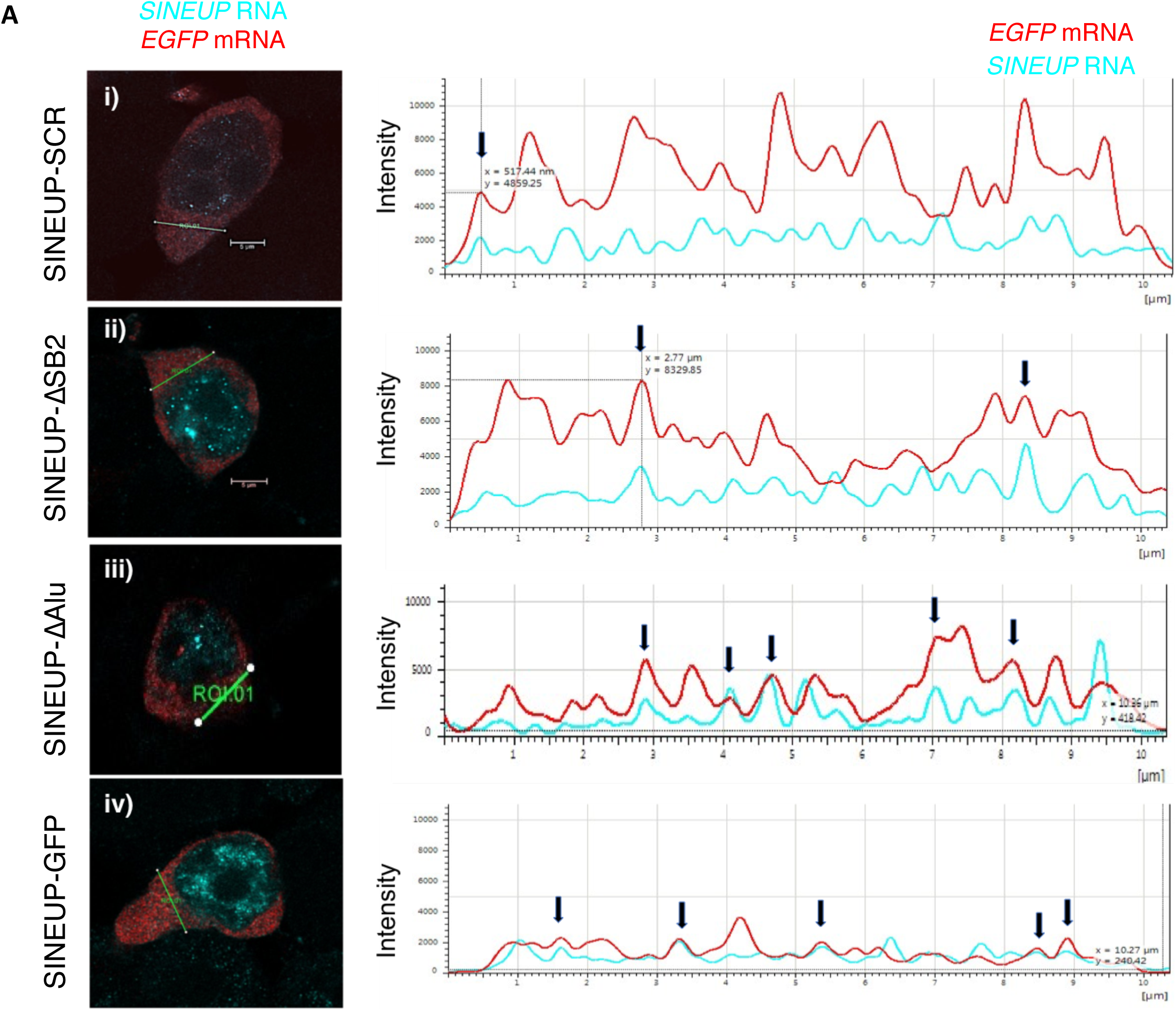
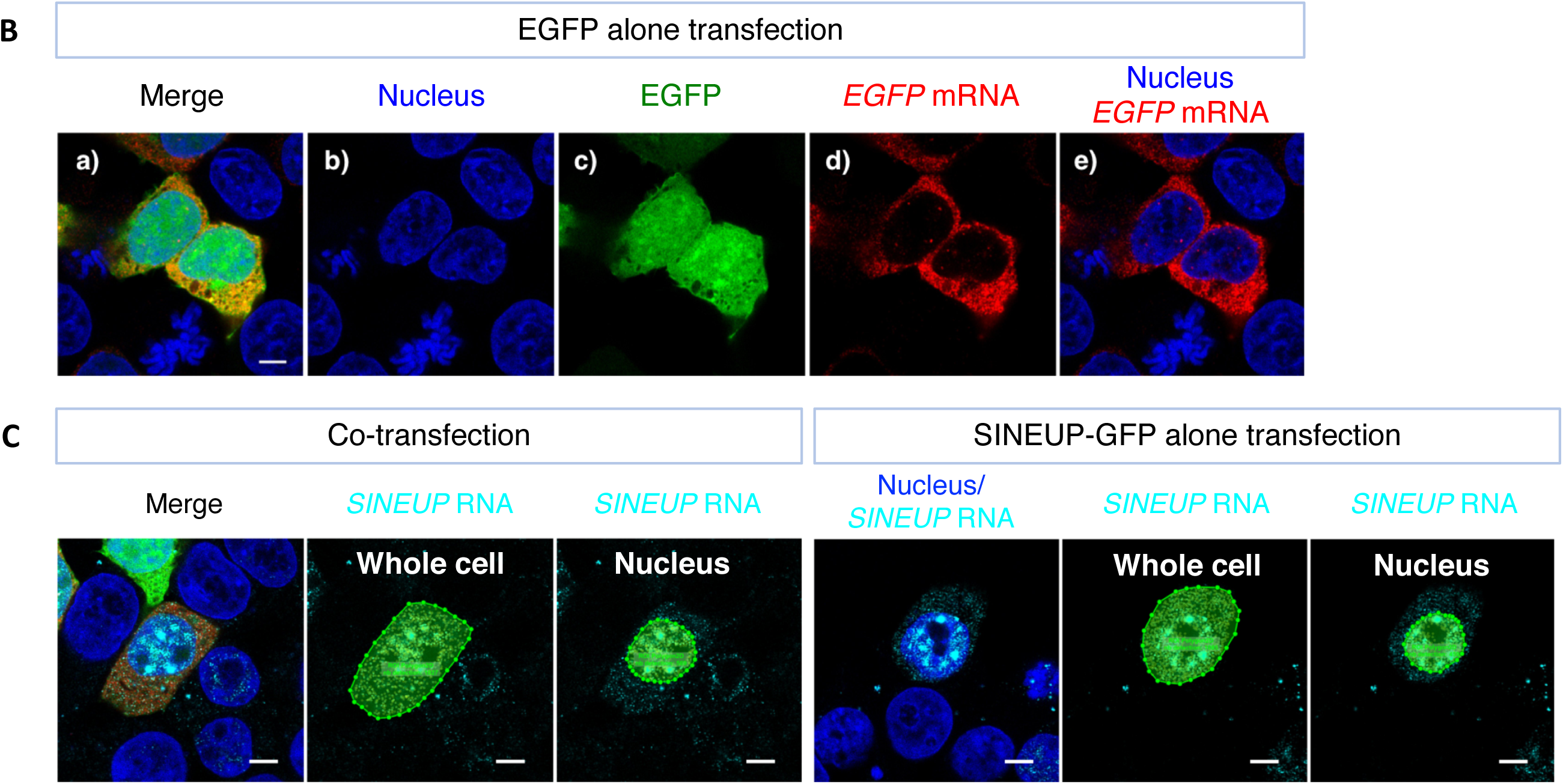
Detection of mutant *SINEUP* RNA signals. A The RNA intensity images. In the graphs on the right, red lines show *EGFP* mRNA and light blue lines show *SINEUP* RNA signal intensity in the region of interest (ROI) indicated as a line between points in the cytoplasm of each image on the left. The X axis shows distance along the ROI line, and the Y axis shows signal intensity. Arrows show the peaks that *EGFP* mRNA and *SINEUP* RNA signals overlape. B Subcellular localization of *EGFP* mRNAs following transfection with EGFP expression vectors alone. Bars indicate 5 μm. C Spot detected images in the cells with co-transfection and with SINEUP-GFP alone transfection. Using icy Spot Detector, signals were detected from the whole cell region and from the nucleus region. Bars indicate 5 μm.

**Fig EV3.**
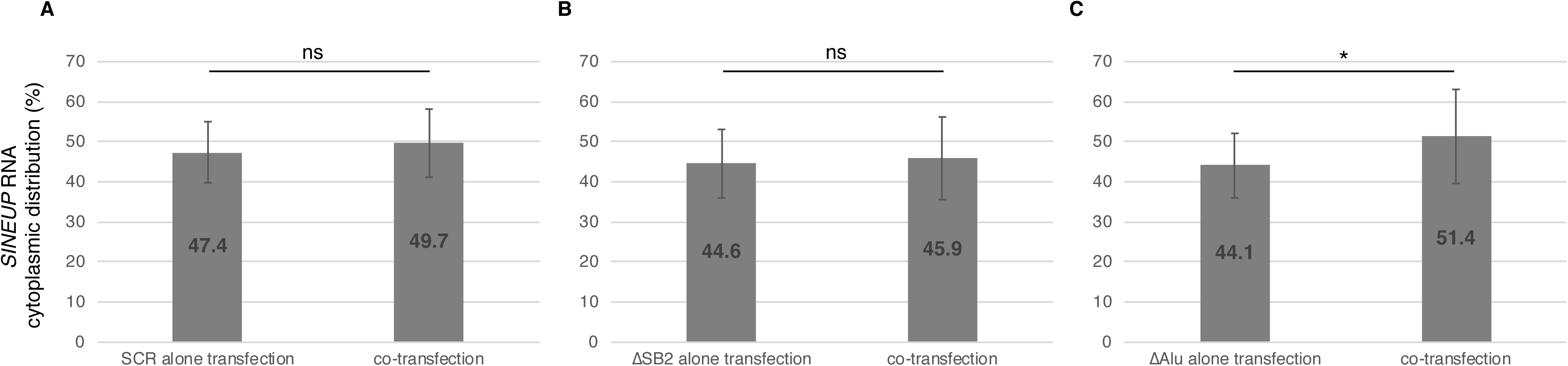
Comparison of subcellular distribution of SINEUP mutants in cells transfected with *SINEUP* RNA vector alone and cells co-transfected with EGFP and *SINEUP* RNA vectors. A, B, C The ratio of cytoplasmic distributions of the SINEUP mutants SINEUP-SCR (A), SINEUP-ΔSB2 (B) and SINEUP-ΔAlu (C) were compared with and without transfection of EGFP vector. *p < 0.05, ns: not significant by Student’s *t*-test. Data are means ± SD of at least 10 independent cell images.

**Fig EV4.**
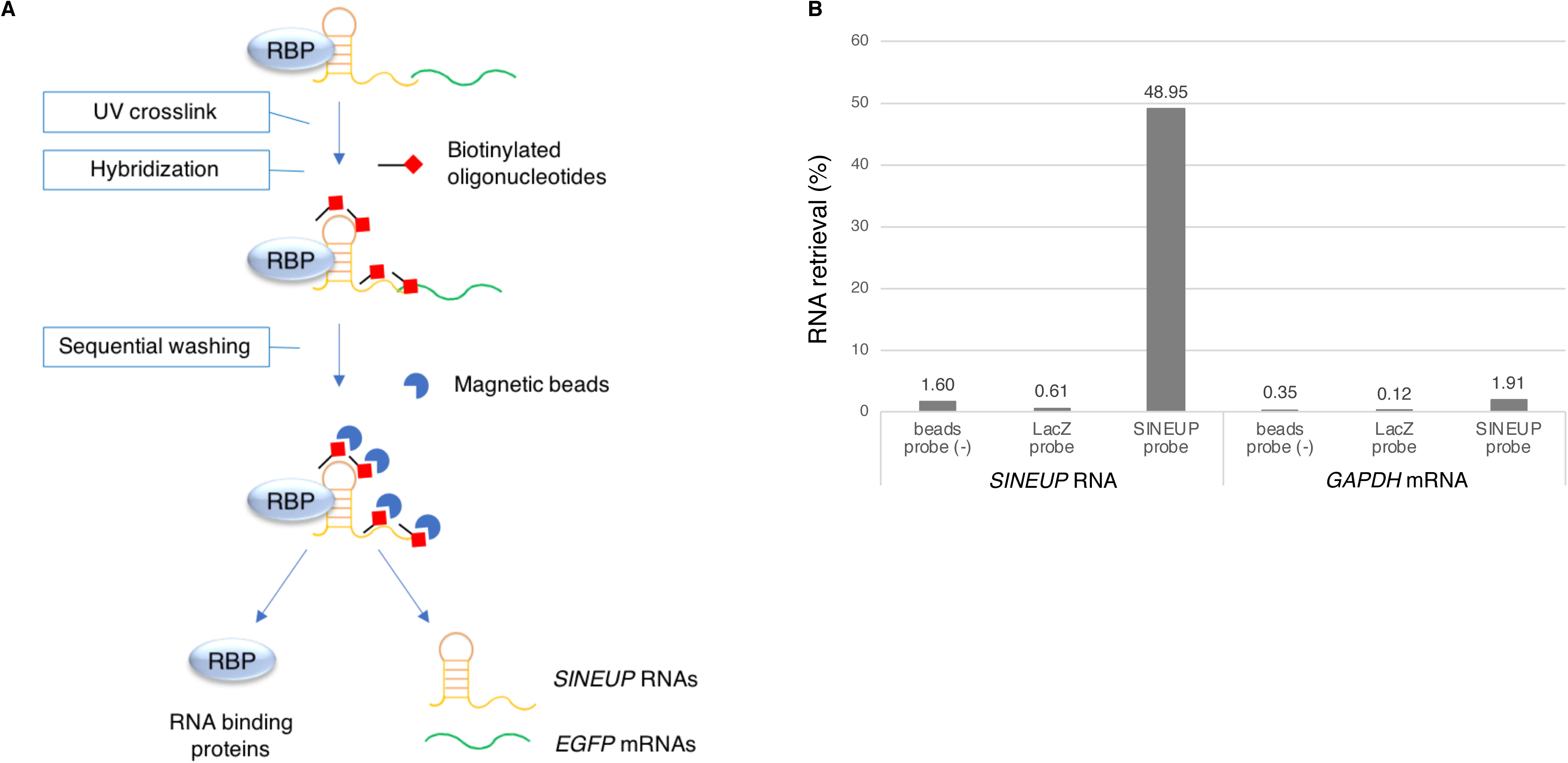
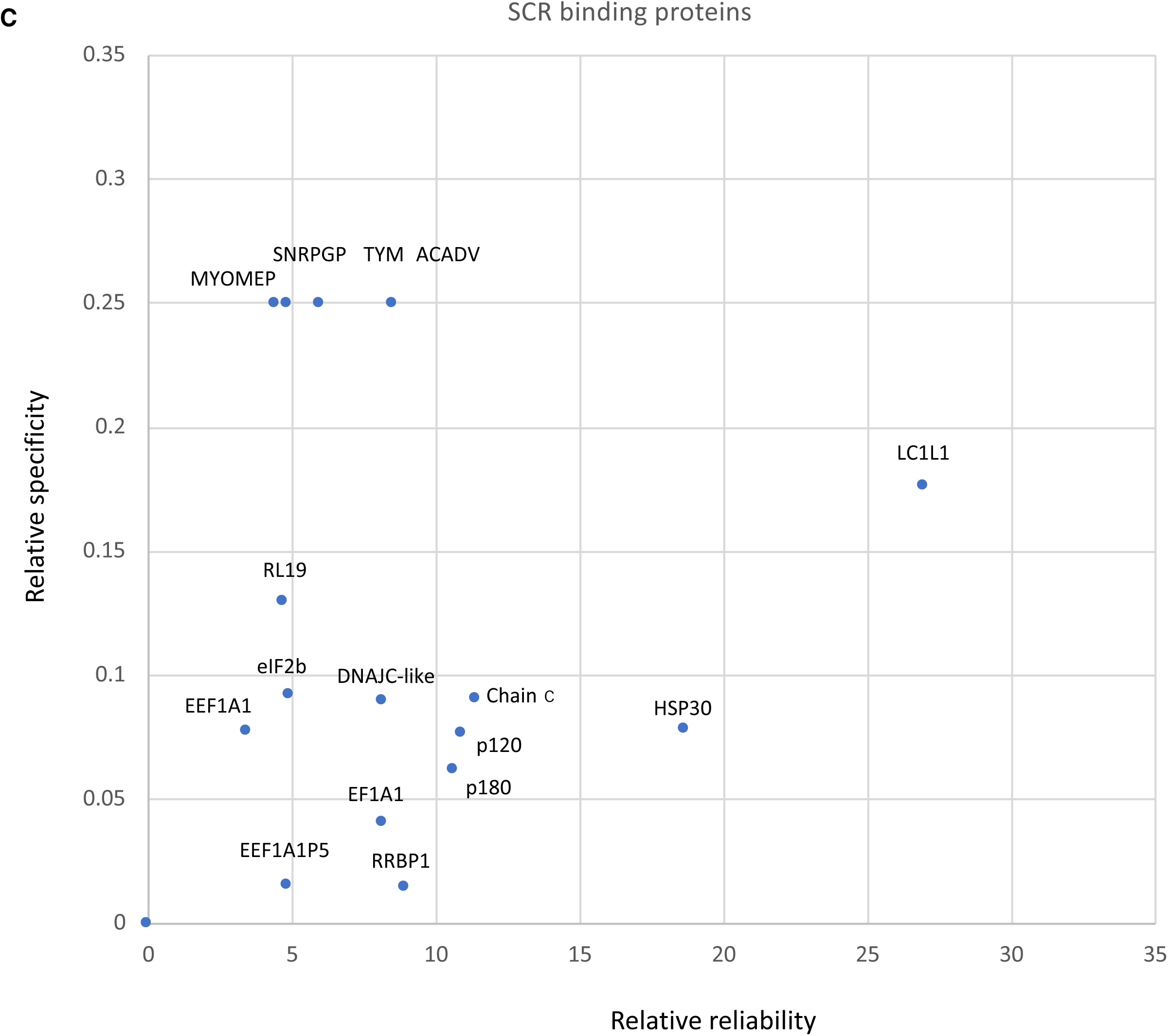
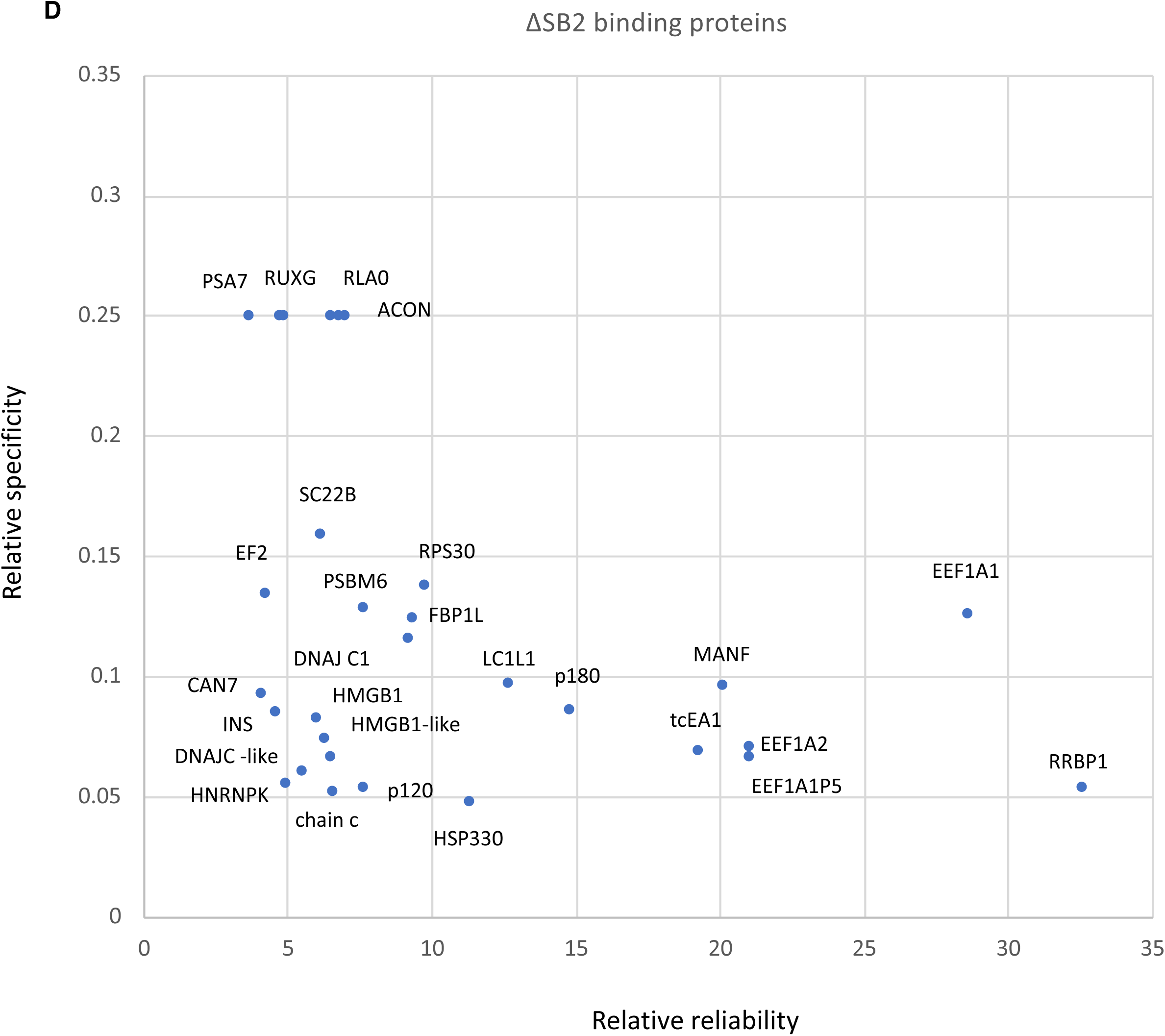
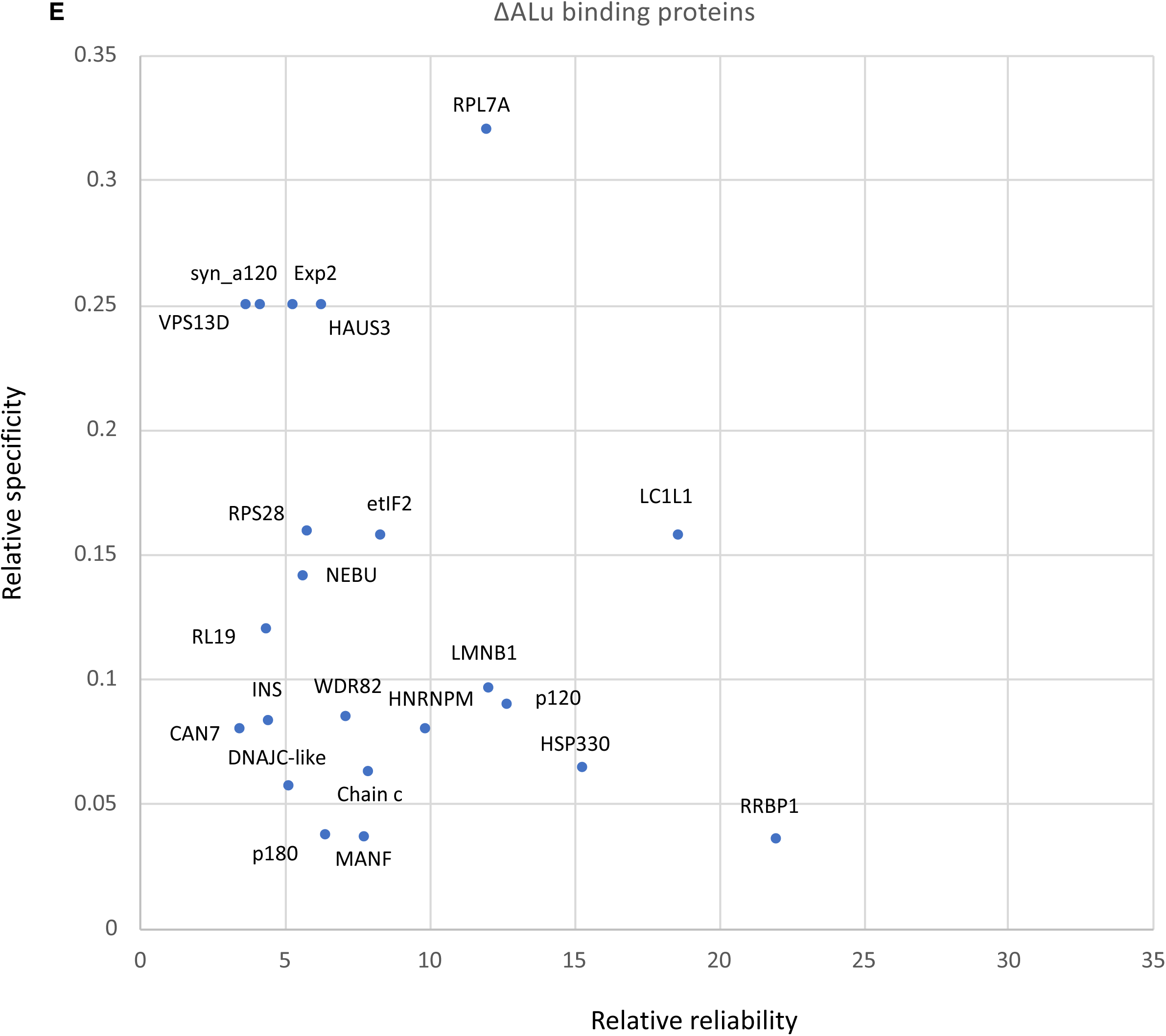
SINEUP RNA-protein interaction. A Schematic workflow of the modified ChIRP protocol (Chu, et al, 2015). B Percentage retrieval of RNA by modified ChIRP with SINEUP RNA probe. Biotinylated SINEUP-GFP probe specifically enriched *SINEUP-GFP* RNA compared with the Magna ChIRP negative control probe (LacZ) and magnetic beads alone (probe (-)). As a negative target RNA, *GAPDH* mRNA was tested to assess non-specific interaction. C, D, E SINEUP-SCR (C), SINEUP-ΔSB2 (D), and SINEUP-ΔAlu (E) RBPs plotted according to relative reliability and specificity.

**Fig EV5.**
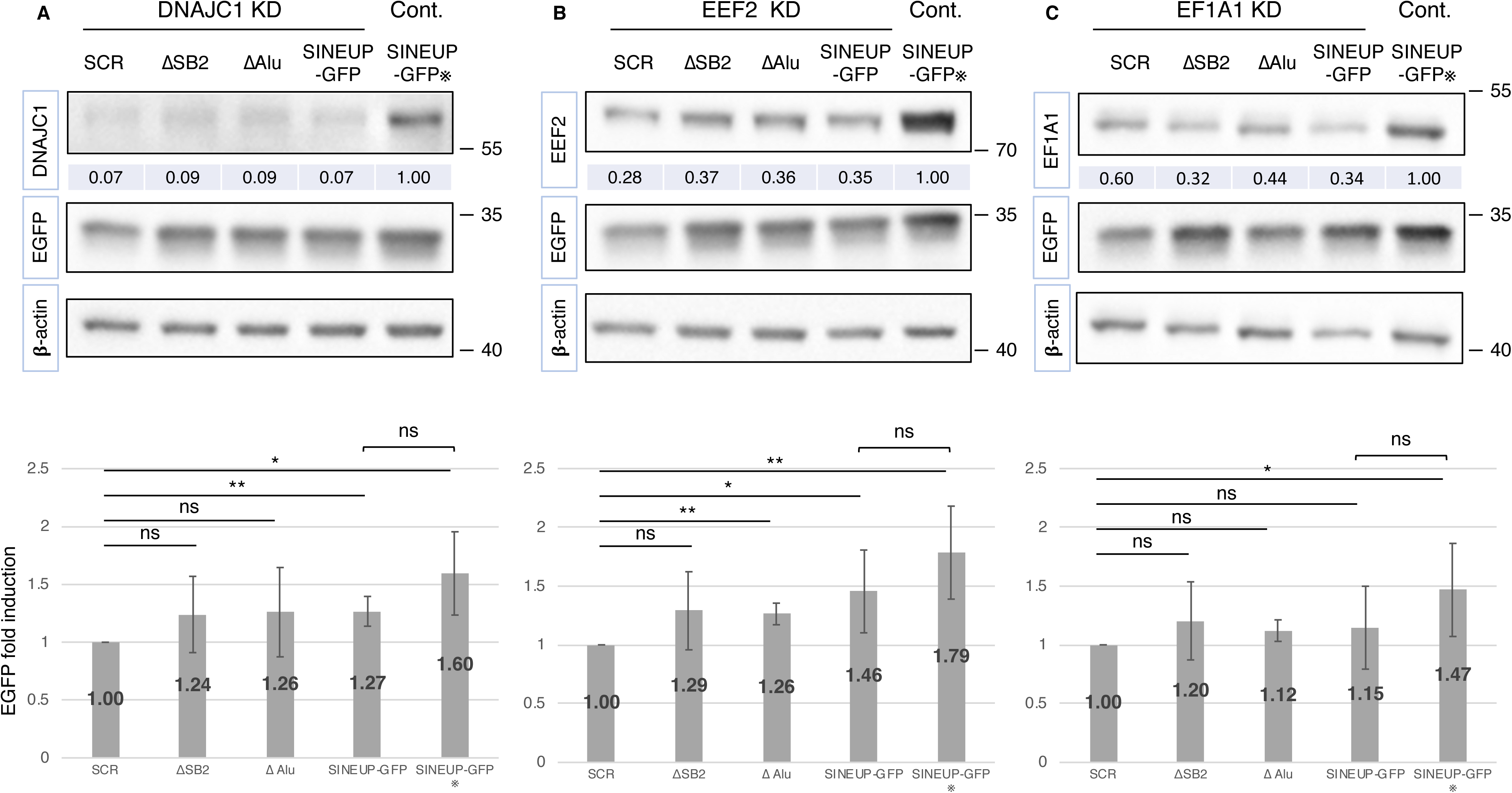
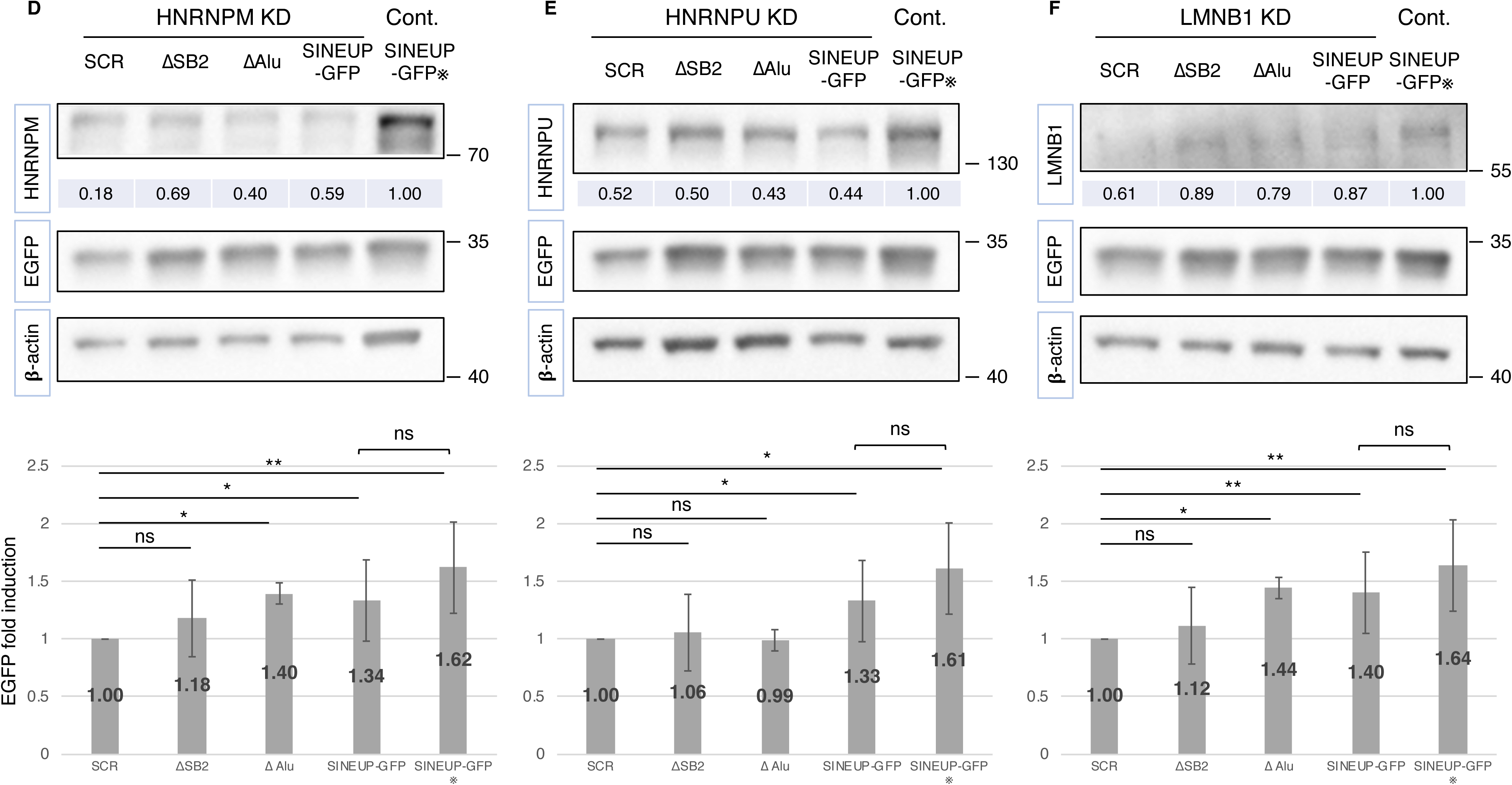
Knockdown of SINEUP RBPs. A–F Knockdown of SINEUP RBPs. Numbers at the top row under the image show knockdown efficiency compared with cells transfected with the SINEUP-GFP vector and negative control siRNA (SINEUP-GFP※ in Fig 4C1 and C2). *p < 0.05, **p < 0.01, ns: not significant by Student’s *t*-test. Data are means ± SD of at least 3 independent experiments.

**Fig EV6.**
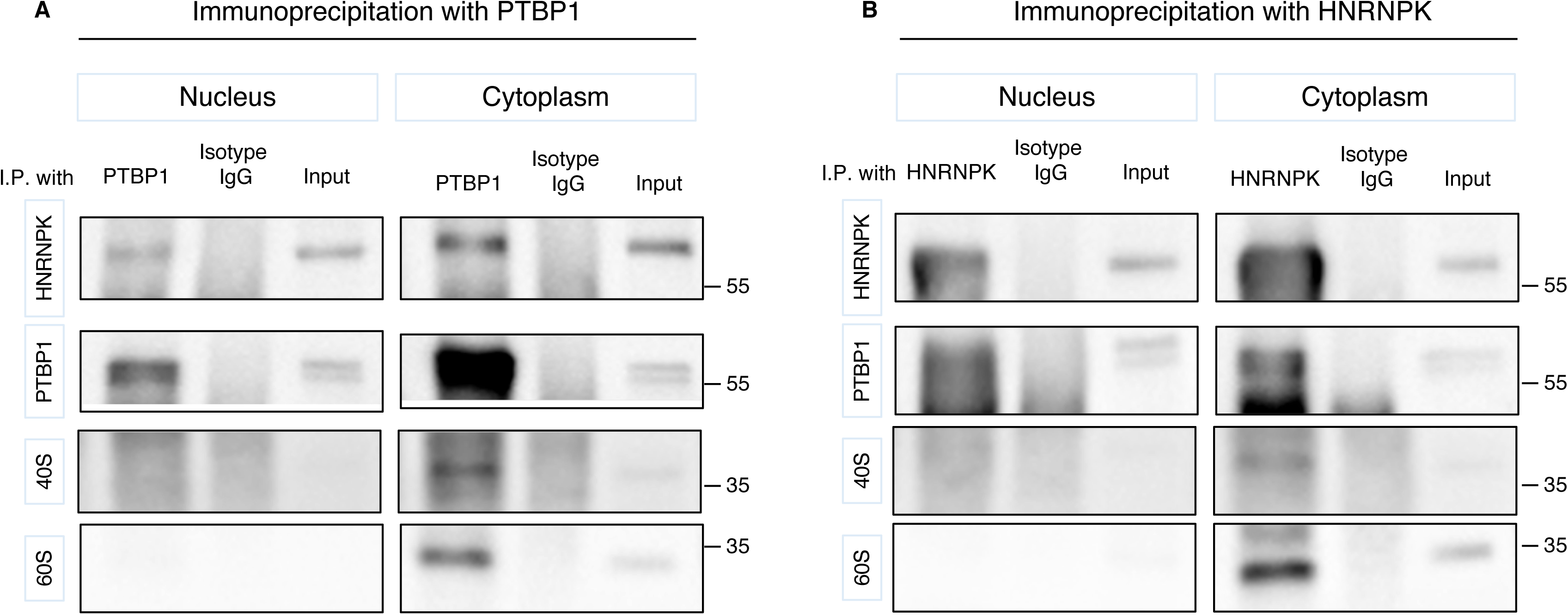
Protein–protein direct interaction by chemical crosslinking with bis (sulfosuccinimidyl) suberate (BS3). A, B Representative Western blot images of immunoprecipitation (I.P.) with PTBP1 antibody (A) or HNRNPK antibody (B). Isotype IgG was utilized as the negative antibody control.

**Table EV1. Mass spectrometric raw data**

Replicates of RBPs with magnetic beads (background), LacZ (negative control), mutants SINEUP-SCR, SINEUP-ΔSB2, SINEUP-ΔAlu and SINEUP-GFP by mass spectrometry are listed.

